# CNN and transfer learning-based classification model for automated cow’s feeding behaviour recognition from accelerometer data

**DOI:** 10.1101/2022.07.03.498612

**Authors:** Victor Bloch, Lilli Frondelius, Claudia Arcidiacono, Massimo Mancino, Matti Pastell

## Abstract

Due to technological developments, wearable sensors for monitoring farm animal behaviour have become cheaper, with longer life and more accessible for small farms and researchers. In this study, an acceleration measuring tag connected by BLE for monitoring behaviour of dairy cows was used. An optimal CNN-based model for the feeding behaviour classification was trained and the training process was analysed considering training dataset and the use of transfer learning. A classifier based on a neural network was trained by acceleration data collected in a research barn. Based on a dataset including 33.7 cow*days (21 cow recorded during 1-3 days) of labelled data and an additional free access dataset with similar acceleration data, a classifier with F1=93.9% was developed. The optimal classification window size was 90s. In addition, the influence of the training dataset size on the classifier accuracy was analysed for different neural networks using the transfer learning technique. During increasing of the training dataset size, the rate of the accuracy improvement decreased, and, starting from a specific point, the use of additional training data can be impractical. Relatively high accuracy was achieved with few training data when the classifier was trained using randomly initialised model weights, and higher accuracy was achieved when transfer learning was used. These findings can be used for estimation of the necessary dataset size for training neural network classifiers intended for other environments and conditions.

**Highlights:** CNN cow feeding behaviour classifier was optimised for neck tags.

Transfer learning technique significantly improves accuracy of CNN.

Relation between the training dataset size and CNN accuracy was estimated.

NN trained by one dataset can be inapplicable for others.

BLE tags measuring acceleration transferred data in real time.

## 1. Introduction

Farm animal activity recognition is important for livestock health and welfare monitoring. Sensors for behaviour recognition of dairy cows have been developed and produced for at least two decades (García et al., 2020). Numerous commercial systems based on acceleration tags (Borchers et. al, 2017; Rayas-Amor et al., 2017; Grinter et al., 2019) provide high accuracy in behaviour recognition. Nevertheless, the commercial systems usually do not provide the access to the raw acceleration data, which is highly important for researchers studying the animal behaviour and developing new methods for efficient farm management. In addition, the price of the equipment and its maintenance is impractical for small farms or farms with middle-sized animals, such as small ruminants.

New sensors are constantly being developed in research. This has been inspired by new technologies providing smaller device size (Liu et al., 2018), better data transfer possibilities, and lower energy consumption, using such as Bluetooth low energy - BLE (Arcidiacono et al., 2021). Due to progress in data processing methods, such as deep machine learning methods, accuracy and robustness of the algorithms monitoring the animal behaviour have been constantly improved. Machine learning model development process includes data pre-processing (e.g., handling the records with missing data, filtering the raw time-series, calculating additional time-series, segmenting the time-series into time-windows), calculating features for some classifiers, and model training and postprocessing. Riaboff et al. (2022) provides an extensive review of these aspect in livestock applications.

In recent years, methods based on Convolutional Neural Networks (CNN) have been widely used for recognition applications, such as human activity recognition (HAR) (Ferrari et al., 2021; Wang J. et al., 2019; Gil-Martin et al., 2020). In livestock applications, CNN for acceleration data measured by tags fitted to animals has been used by Pavlovic et al. (2021), Peng et. al (2019) and Li, C. et al. (2021). Design of a NN-based classifier includes a number of factors, particularly, the type and the architecture of the NN must be fitted to its application. For livestock, Convolutional NN (CNN) and Recurrent NN (RNN) with 2-4 convolutional layers have been utilised to process time series data.

To train deep neural networks effectively, a big amount of reference data must be usually collected and labelled (Russakovsky et al., 2015). Different sensors are used to achieve labelled reference for the cow behaviour and body position: e.g., feeders for estimating the feeding time (Pastell et al., 2018) or halter to measure the rumination and feeding time (Pavlovic et al., 2021). Manual labelling can be done from direct cow observations (Riaboff et al., 2020) or video recorded by cameras installed in the cow environment (Arcidiacono et al., 2017), however, this method is highly time consuming. In cases where the observations on actual behavior cannot be done, unsupervised methods for the behaviour classification are used (Shahriar et al., 2016). To advance the development of HAR, some researchers published data used for their studies in open access, saving time for reference preparation and enabling the use of larger datasets for model training (Wireless Sensors Data Mining - WISDM). According to Riaboff et al. (2022), a large variety of collected data for the classifier training have been used in different studies (from 2 to 200 hours), and a recommendation to collect at least from 25 animals for at least 40 hours was given. However, no analysis of a required amount of the training data was found for farm animal activity recognition and HAR.

Different types of data augmentation have been used to enlarge the training dataset (Kalouris et al., 2019; Li at al., 2021): rotation, permutation, jittering and scaling performed for the original signal, or local averaging as a down-sampling technique and shuffling in the feature space (Eyobu et al., 2018). Optimisation of the classification window length (epoch, time-window, segment, observation) was done in each study for its specific datasets.

Transfer learning is a method that prepares a classification model for one problem type and uses this pretrained model as a base for training a model for other but similar problem type; for example, training a pretrained model based on younger population groups and using it as the initial condition to train a model for older people, as was done by Kalouris et al. (2019). Additional training of pretrained models by data from specific objects and environment improves their accuracy and decreases the training time relative to newly trained models or existing models. In HAR, this method was used by Oh et al. (2021), Li, F. et al. (2020). According to our review, this method has not been used for livestock activity recognition.

The aims of this study were to find an optimal behaviour classification model and evaluate the impact of the size of training dataset and the use of transfer learning on the accuracy of farm animal activity recognition classification using CNNs. We used a low-cost open-source system based on acceleration tags to develop a behaviour classification method according to the best practices taken from the reviewed studies. We evaluated the minimal amount of data needed for effective training of a NN classifier and the use of transfer learning based on openly available dataset.

## 2. Materials and Methods

### 2.1. Barn study area and monitored cows

The data used for development and validation of the system was collected from dairy cows housed in a free stall research barn (Luke, Maaninka, Finland). The barn comprised two separated compartments with a 10×20 m area containing, in total, 48 lying stalls and 24 feeders (Hokofarm, The Netherlands). A group of 48 cows, specifically Ayrshire (n=18) and Holstein (n=30), were housed in the study area during the lactation period. The barn was equipped with continuously recording cameras (HDBW5541R, 5M, 3.6mm, Dahua, China) installed on the ceiling and covering the major part of the area.

### 2.2. System design

The tags measuring 3D-acceleration (RuuviTag, Ruuvi Innovations, Finland) were packed in plastic boxes and adjusted by a Velcro belt on the left side of the cow collars or to one of the legs just above the metatarsal joint (Fig. 1). In total 96 tags were fitted to the collars and legs of 48 cows. The tags attached to the legs were used only to test the reliability of wireless transmission with increased number of tags. The tags broadcasted the data as BLE advertising packets. The acceleration was sampled at 25Hz, and the frequency of the message sending was 5Hz. Each packet included five samples for three axes, which amounted to 15 acceleration values. The data from the tags were received by 6 receiving stations, which were single-board computers equipped with a Bluetooth antenna (Raspberry Pi 3 B+, Raspberry Pi Foundation, UK). The stations were packed in hermetic cases with heat sink ribs and installed at 3-5 meters height on the barn structures (Fig. 1 (c)) and evenly distributed in the study area to minimise the distance from the tags. The receiving stations recorded the tag accelerations and the receiving time. The data was stored on the base station’s storage and was sent via a local network maintained by a router (EA7500, Linksys) through a message queuing protocol (ZeroMQ, iMatix Corporation). A PC (Intel(R) Core(TM) i7-9750H, CPU 2.6GHz, RAM 16GB) received the messages and stored the raw data in CSV files. The tag and base station software were written in C++, and C# language was used (Microsoft, USA) for the PC. The lifetime of the tag battery was estimated using power profiling described in Bloch et. al (2020) as about three years.

**Fig. 1.**
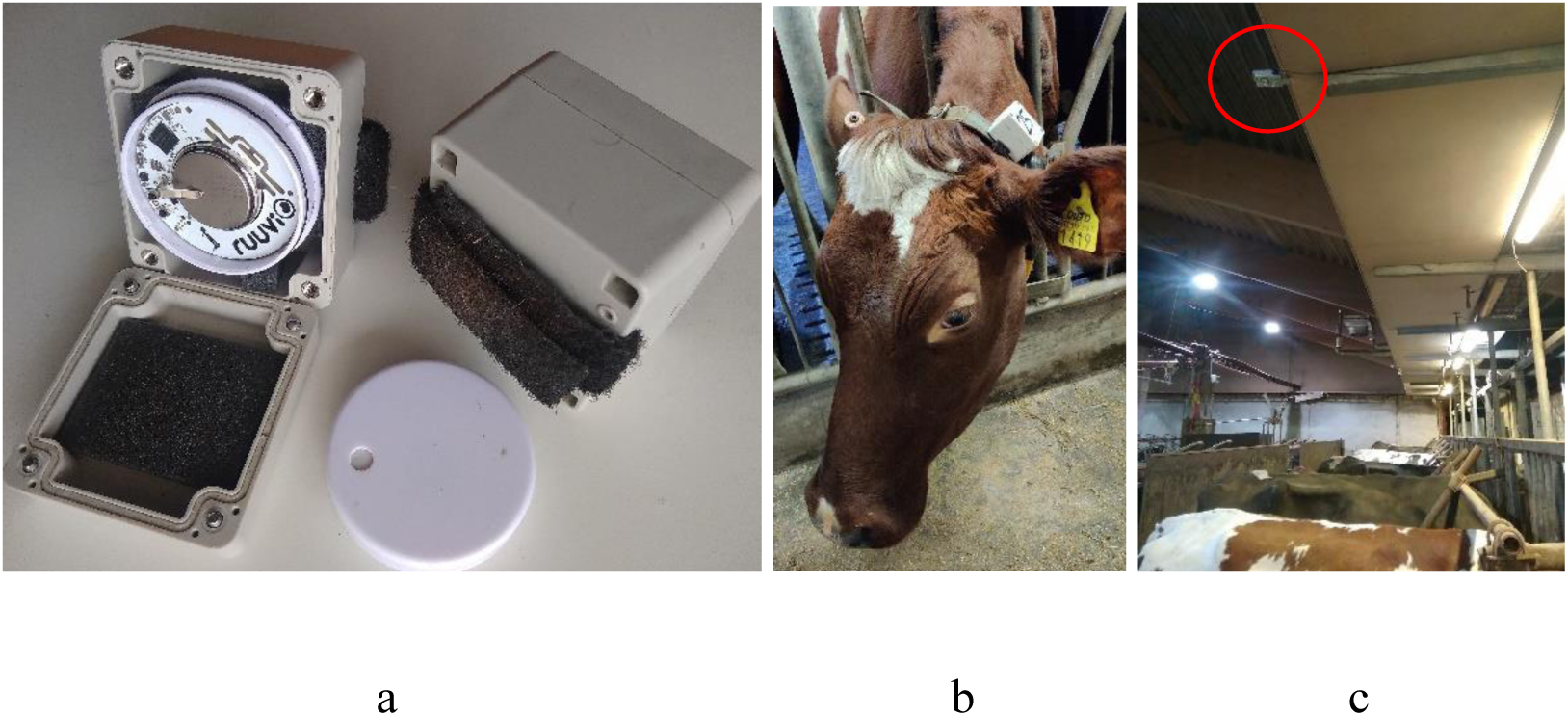
Component of the location and acceleration measuring system installed in a barn: RuuviTag inside a protecting plastic box (a), tag on the cow collar (b), receiving station installed on a barn structure (c).

**Fig. 2.**
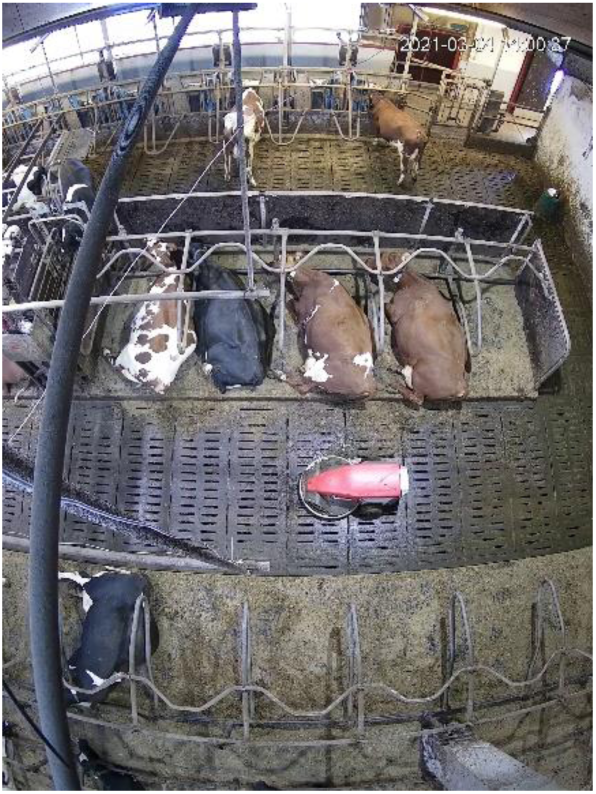
Top-view image of the research barn acquired by one of the cameras.

### 2.3. Data collection and labelling

Three feeding behaviour classes were considered in this study: feeding, ruminating and other (neither ruminating nor feeding).

The individual feeders were used to collect reference data for the feeding. We assumed that animals were not ruminating while registered to the feeders. Manual labelling was used to recognise ruminating and other behaviour. Because of restrictions in the camera views, only time intervals, where the behaviour was clearly detected, were labelled. Individual cows were recognised by their unique colour patterns of coat; images of cows from both sides, on top and from the head were captured at the beginning of the experiment to aid the recognition. The time labels of the feeders and the cameras were synchronised with the time label of the tag receiving stations with an accuracy of 1 sec.

The average rate of the missing data messages was 52.6±6.1% (mean±STD). The missing samples were concentrated in groups containing multiplication of five samples (since the data was transferred in packets including five samples) as shown in Fig. 3.

**Fig. 3.**
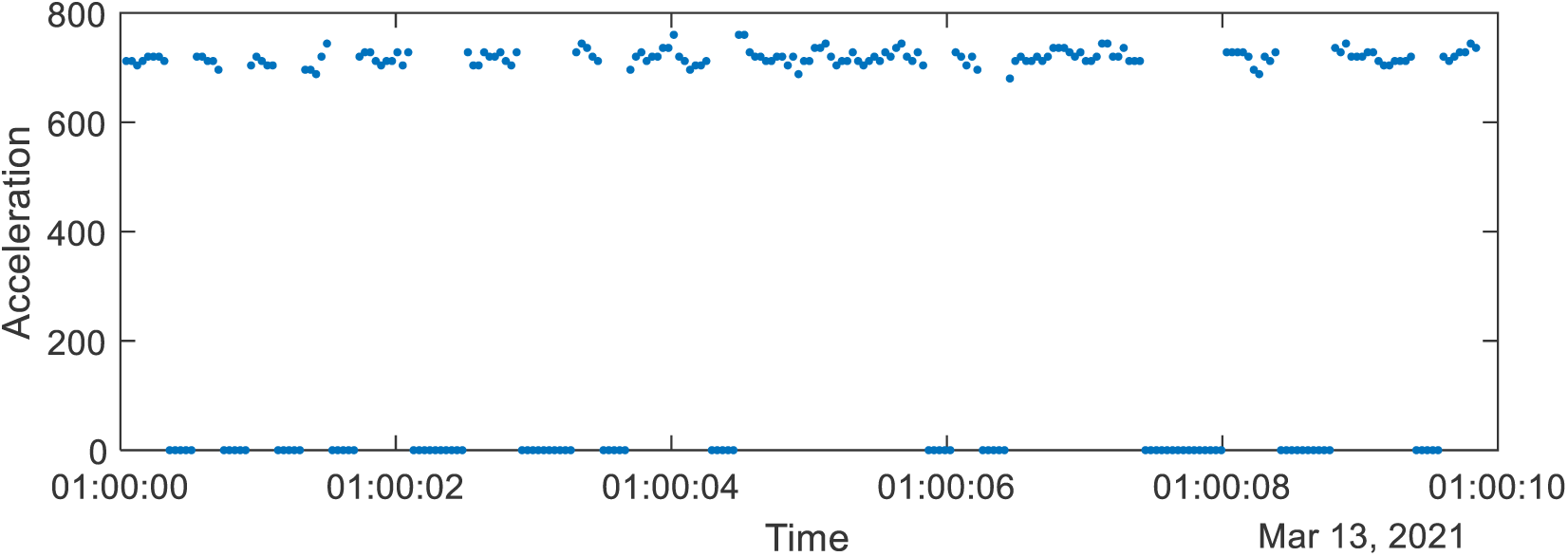
Illustration of the missing samples in the recorded acceleration data.

The classification models were trained on two datasets (Table 1): data collected in this study as explained above (4.3.2021-13.3.2021) and labelled for 21 cows, and an open-source data published by Pavlovic et al. (2021). Differently from the current study, Pavlovic et al. (2021) used sampling at 10Hz and sensors with ability to download the data, thus preventing data loss. All data labelling was done automatically with the help of halters measuring cow feeding behaviour (Rumiwatch, Switzerland).

**Table 1.** Characteristics of datasets used for NN model training (mean±STD).

|  | Number<br>of cows | Period,<br>days | Average time<br>for a cow, hours | Total time,<br>hours (days) | Fe,<br>% | Ru,<br>% | Oth,<br>% |
| --- | --- | --- | --- | --- | --- | --- | --- |
| Collected<br>data | 21 | 1-3 | 38.5±12.4 | 809 (33.7) | 19.7<br>±5.7 | 36.9<br>±6.1 | 43.3<br>±6.9 |
| Open-source data | 18 | 6-18 | 191.7±87.5 | 3450.5<br>(143.7) | 17.6<br>±3.8 | 38.4<br>±3.5 | 43.9<br>±6.6 |

Fe – feeding, Ru – ruminating, Oth – other behaviours.

### 2.4. Data processing

Data pre-processing included filtering, amplitude normalisation, sampling frequency normalisation, augmentation and balancing. The raw acceleration data was filtered by a hamming high-pass filter with filter order 511 and cut-off frequency of 0.1Hz. The acceleration values were normalised to ±1. Because of a high rate of missing samples, the missing data was replaced by zeros to preserve the structure of the time series and used for the training according to the methods proposed by Hossain et al. (2020).

Window overlap augmentation of consecutive windows with 50% overlapping was used to increase the amount of the training data, according to recommendations of Riaboff et al. (2022) and review by Ferrari et al. (2021). Since tags fitted on collars were able to rotate around the neck, to simulate possible rotation around the X axis parallel to the cows’ neck and train the model to be insensitive to the tag orientation, the rotational augmentation was used. The Y and Z acceleration components were rotated in 3D space by the transformation

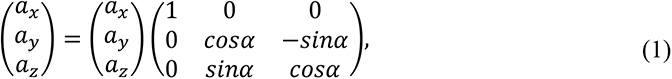

where *a_x_*, *a_y_* and *a_z_* are the components of the tag acceleration measured along X, Y and Z axis, *α* is a random rotational angle, as it was done by several authors (Um et al., 2017; Kalouris et al., 2019; Li at al., 2021). Every classification window was rotated by a random angle.

To balance data for all behavioural classes, data windows from the minor classes were randomly taken, rotational augmentation on a random angle was performed and the result was added to the training set. The balancing was performed for data collected from each individual cow during one day. Increasing of the dataset size because of the balancing depended on the level of imbalance and was on average 41±20%.

A postprocessing of the classified behaviour was done by median filter with the window length equal to five. Both data collected for this study and published by Pavlovic et al. (2021) were processed using the same procedure. Additionally, the sampling frequency of Pavlovic dataset was changed from original 10Hz to 25Hz used in the current system by zero padding.

### 2.5. Tested classifying models

Two NN classifiers found in the reviewed literature were compared in this study:

- CNN2. Methods for the Human Activity Recognition described in Brownlee (2016). The CNN2 consists of two 1D convolutional layers with kernel size 3, dropout layer and pooling layer as presented in Fig. 4.

**Fig. 4.**
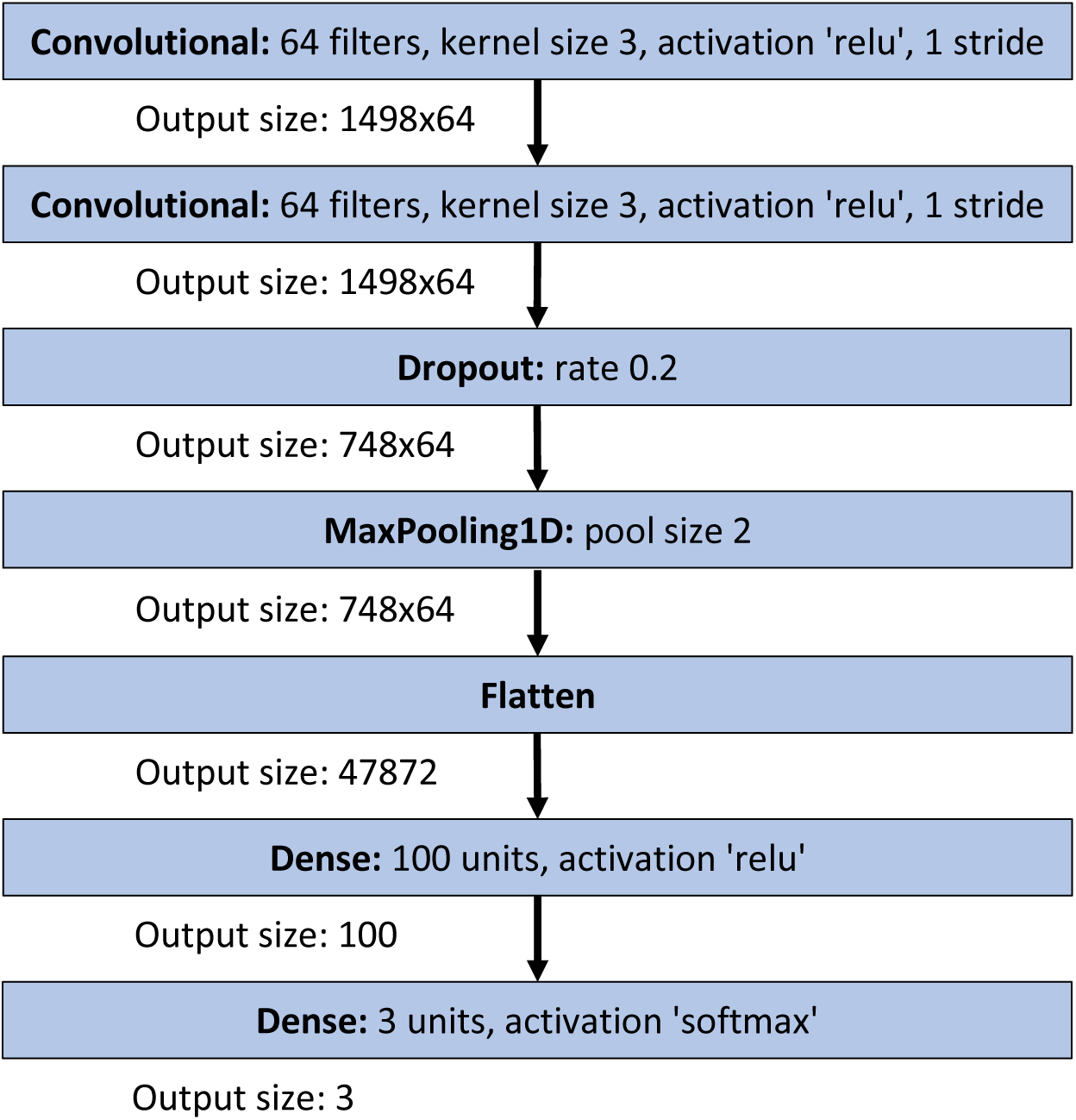
CNN2 architecture.

- CNN4. Deep CNN for cow activity recognition described in Pavlovic et al. (2021). The CNN4 consists of four 1D convolutional layers with kernel sizes 52 and 1, dropout layer and pooling layer.

The structure of the models is available in the supplementary materials https://github.com/cowbhave/CowBhave_BehaviorModel.

The size of the classification window was optimised similarly to what found in the reviewed studies (Table A1: Wang Y. et al., 2019; Riaboff et al., 2022). In this study, the optimal classification window size was searched by a grid search algorithm in the set [5 10 30 60 90 120 180 300]sec with the extreme values 5s and 300s taken from Table A1 for cows. The total amount of the bouts for each behaviour class in the available labelled dataset with a length less than 300s was 3.1% for feeding, 2.7% for rumination and 9% for the other behaviours.

The pretrained classification models for the transfer learning for different window sizes were trained by the data published by Pavlovic et al. (2021). The transfer learning was done by training the last convolutional layer in the models (the second for the CNN2 and the fourth for the CNN4 model) and all subsequent layers. The training was performed for 30 epochs with a 0.001 learning rate using Adam optimiser. The model was implemented and trained using Keras library with Tensorflow.

The datasets used for training (which were the dataset collected in this study and the dataset published by Pavlovic et al. (2021)) were achieved from different cows and environments with different sensors, sampling rate and rate of missing data. Hence, they were used to test the applicability of the trained models for other cows and environments. The classification accuracy of models trained on one dataset and validated by another dataset was estimated.

### 2.6. Analysis on the effect of training dataset size

To evaluate the dependence of the model accuracy on the amount of data used for the model training, the learning using randomly initialized model weights and the transfer learning were performed using different parts of the original dataset. The smallest data amount, which can be used for training, is the classification window. For this test, only 60s window, including 60*25=1500 acceleration samples (where 25Hz was the accelerometer sampling frequency) was used. Each window had its behaviour class label, hence, it was defined as a training sample and the dataset size was measured by the number of training samples. Small testing datasets were created as parts of the original dataset. The original acceleration data for one cow during one day was stored in one file, resulting totally in 56 files. To create the minimal dataset, one training sample for all three classes was taken from each file, totally 3*56=168 training samples (3*56*60s=10080s=2.8hours). This amount represents 0.34% of the total dataset size of 48540 training samples. For the next dataset including two training samples taken from each file, the dataset size was 3*56*2=336 training samples. Hence, in the analysis, the initial datasets including the following dataset sizes were used: 168, 336…1680, 4854, 9708…48540. In bigger datasets, where the equal amount of training samples for each behaviour class could not be taken, actual data size for the training was increased by the balancing. The sizes of the corresponding datasets enlarged by the augmentation and balancing were: 336, 504…3192, 13878, 28358…132762. For each fold in the 10-fold validation, the size of dataset was multiplied by a factor equal to about 9/10.

### 2.7. Accuracy evaluation

The accuracy was estimated by total classification precision and macro F1, confusion matrix, and (micro) *Precision_i_*, *Recall_i_* and *F*1*_i_* for each separate behaviour class *i* as follows:

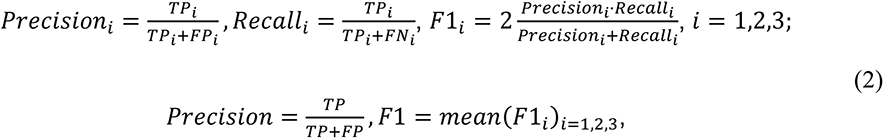

where *TP_i_* is the number of true positive classifications for the class *i*, *FP_i_* is the number of false positive classifications for the class *i*, and *FN_i_* is the number of false negative classifications for the class *i*. TP is the total number of correct classifications for all classes, FP is the total number of incorrect classifications for all classes.

## 3. Results

Performance of the CNN2 model for the 60s window size, trained using random weights and by transfer learning for different dataset sizes used for the training, is presented in Fig. 5. The use of transfer learning clearly improved the performance (F1 score) of the classifier for small dataset sizes and F1-scores 8x-9x% were obtained with just 336 training samples. The advantage of using transfer learning disappeared when over 24 000 training samples were used.

**Fig. 5.**
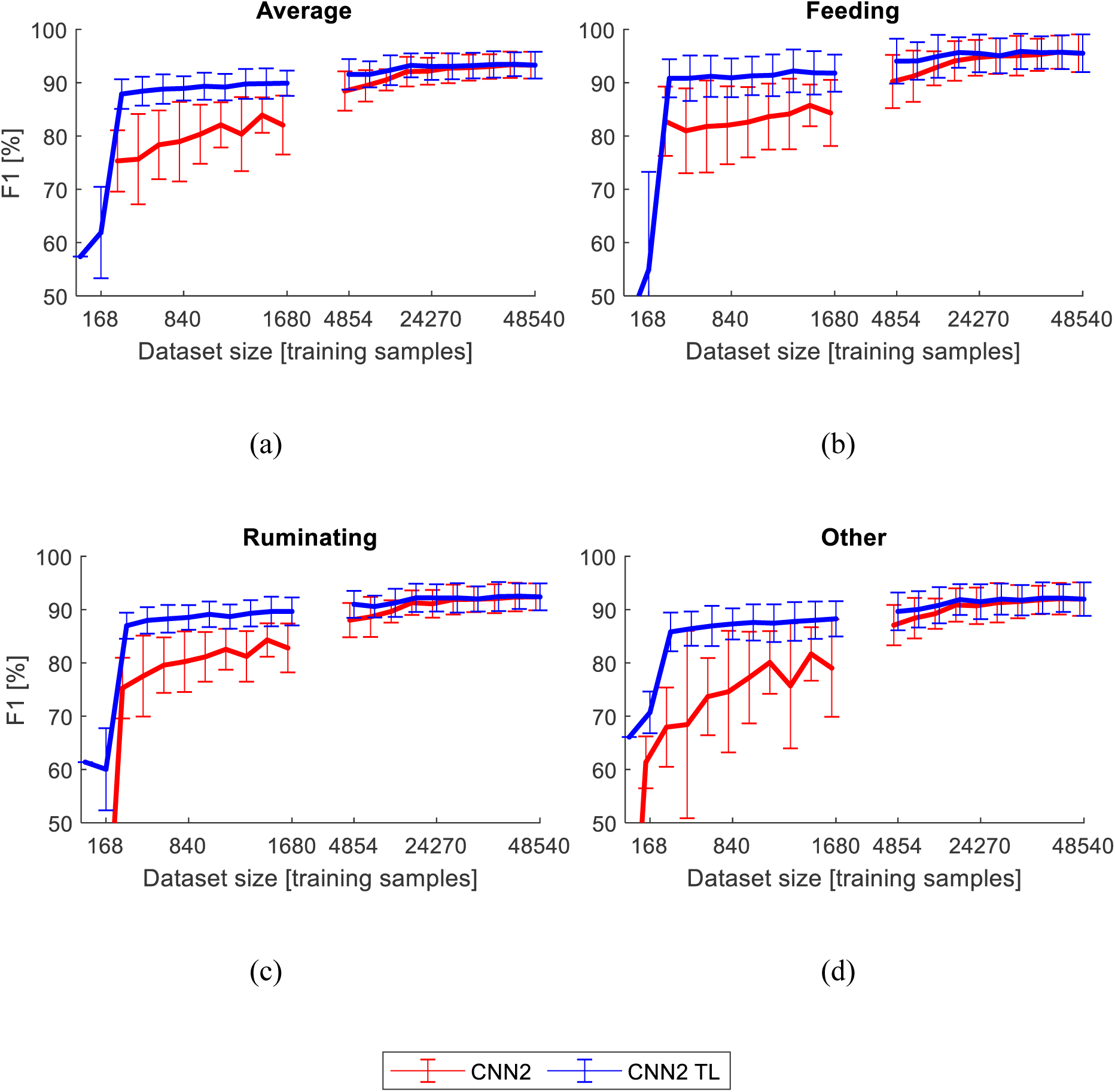
F1 score of the model CNN2 trained using randomly initialized model weights (CNN2) and by the transfer learning (CNN2 TL) for the window sizes 60s depending on the training dataset size measured in training samples taken from the original dataset. The corresponding actual data after augmentation and balancing is: 336,..,3192, 13878,..,132762. The error bars represent the STD for 10-fold validation.

The training window size had a clear effect on model performance when training the models on the whole original dataset. F1 score for the classifiers CNN2 and CNN4, trained using randomly initialized model weights and by the transfer learning for all tested window sizes, is presented in Fig. 6. The highest F1-scores were obtained for a 90s window size.

**Fig. 6.**
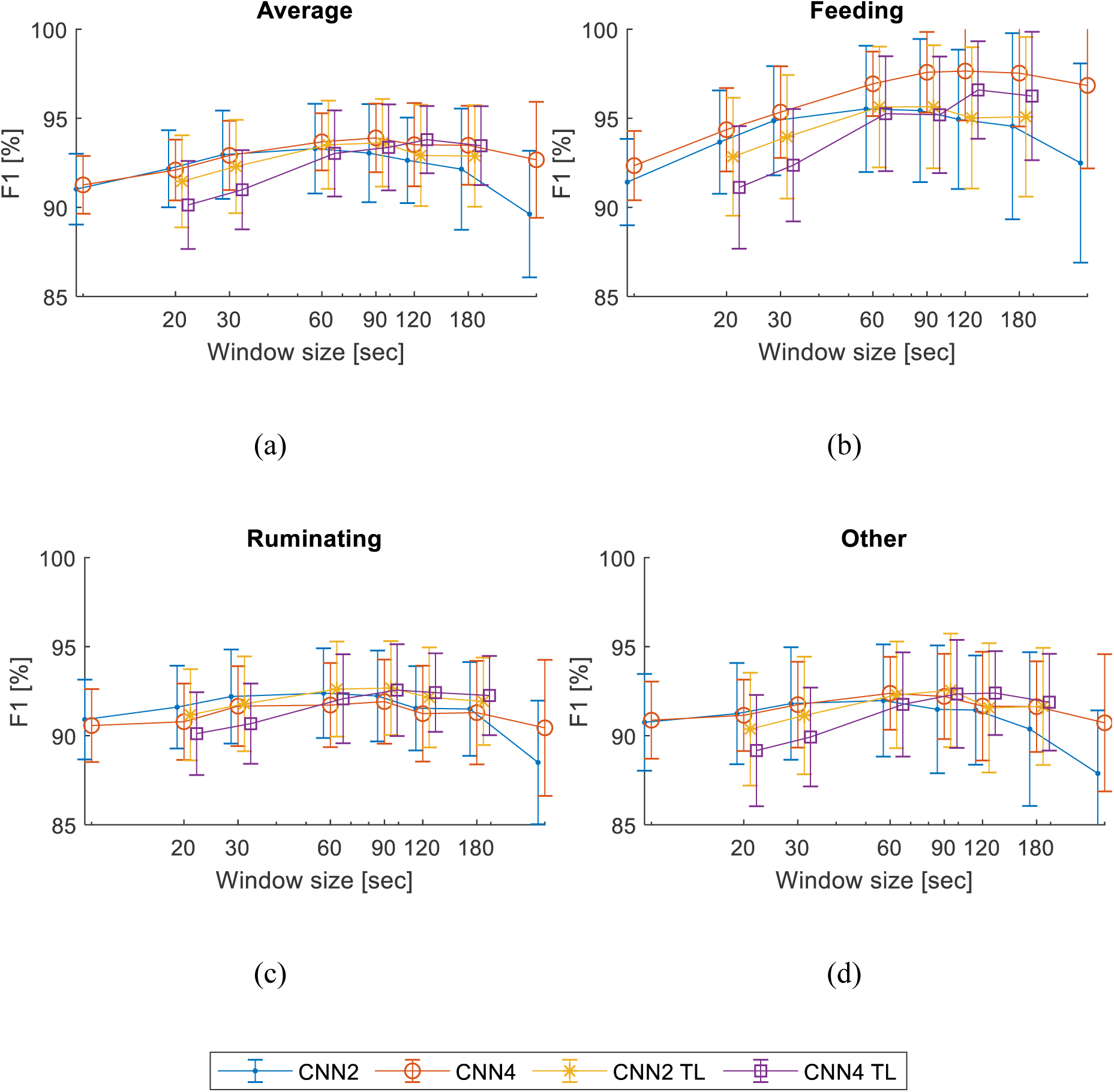
Performance of the tested models CNN2 and CNN4 trained using randomly initialized model weights and by the transfer learning (CNN2 TL and CNN4 TL) depending on the window size. The error bars represent the STD for 10-fold validation.

The best accuracy scores of all classifiers trained with the full dataset for window size with the highest average F1 score for each model are presented in Table 2. There were only minor differences in performance between the simpler CNN2 and more complex CNN4, and between using transfer learning and training using randomly initialized model weights. For all the tested models, clear optima were in the 60-120s range.

**Table 2.**
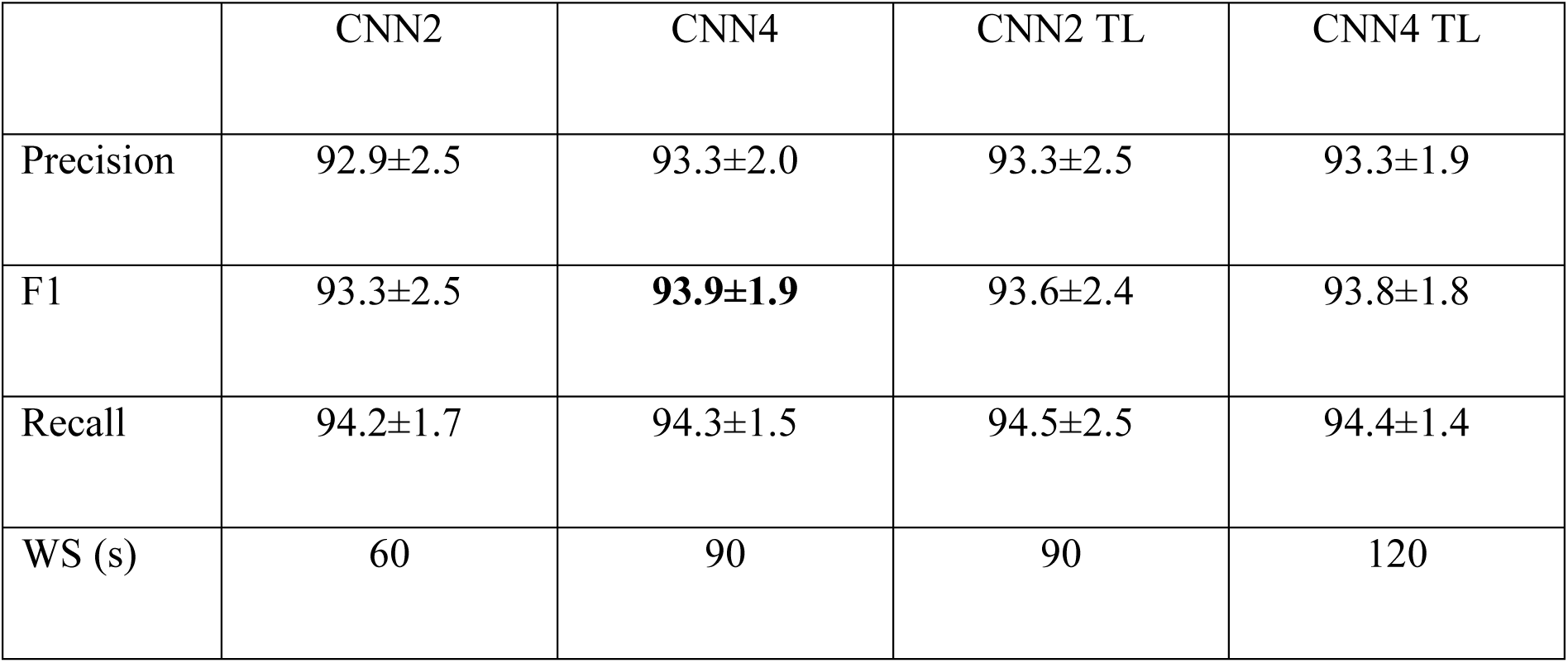
Comparison of model performance trained by simple training and transfer learning. Precision and F1 values are given averaged (mean±STD) for 10-fold validation for optimal window sizes (WS).

The accuracy of the models trained on one dataset and validated by another was low: F1=57.3% for the model trained by the dataset published by Pavlovic et al. (2021) (the pretrained model before transfer learning), and F1=63.0% for the model trained by the dataset collected in this study and validated by dataset published by Pavlovic et al. (2021).

## 4. Discussion

Using the transfer learning technique, relatively high (F1 = 87%) behavioural classification accuracy was reached with less than 500 training samples and a significantly better performance (12% higher) compared to model initialised with random weights. Even higher classification performance was reached by using more training data. In this study, the average F1-score reached its maximum level with around 30 000 training samples and the use of transfer learning was beneficial up to 24 000 training samples.

The differences in accuracy between simpler CNN2 and more complex CNN4 architectures and between learning model using randomly initialized model weights and transfer learning were not significant when the full dataset was used. It may suggest that increasing the number of layers in a NN does not significantly increase the classifier accuracy for accelerometer data. Additionally, using transfer learning can improve the model performance when large amount of training data cannot be collected.

The analysis of different training dataset sizes showed a constantly increasing accuracy when more training data is used. However, it shows the limitation of the classifier as well, since increasing the dataset size to over 30% does not significantly increase the F1 score from the level of approximately 94%. Nevertheless, this effect occurred in this specific case where the data was collected with specific sensors in the same environment for the same animals during a short period of time. The influence of the condition diversity on the accuracy of the models should be further studied.

A feeding behaviour classification model with the best performance was CNN4 with an average 93.9% F1 score for the 90s optimal window size, which is close to the median among the values found in the literature review (Table A1). In practise, classification models are run on continuously measured data which is not split into windows containing data from only one behaviour. Large window size can therefore create additional uncertainty in the classification accuracy because sample windows can contain mixed behavioural classes. The F1 score achieved in this study was high compared to the systems using NN for cow behaviour classification such as those by Pavlovic et al. (2020) with F1=82%, Peng et al. (2019) with F1=88.7% and Li, C. et al. (2021) with F1=94.4%. Among the systems using machine learning with the performance reviewed in Table A1 were those by Arcidiacono et al. (2017) with F1=93.3%, by Tian et al. (2021) with F1= 98.51%, and by Riaboff et al. (2020) with a total accuracy of 98%. Exact comparison of the accuracy scores is impractical because of differences in the research conditions, such as experiment environment, sensors and amount and type of the collected data. However, the results in the mentioned studies can be applicable in actual conditions. Having the data openly available would be beneficial for comparison of developed methods. Data collected in this study and other freely available datasets are listed in https://github.com/Animal-Data-Inventory/PLFDataInventory.

The low accuracy (F1=57.3% and F1=63.0%) of models trained and validated by datasets achieved from different cows, different sensors, different reference data and environments showed that both models were not applicable for other environments without additional fitting. However, the high accuracy of the transfer learned models can mean that the basic patterns of the cow motions characterising this class of problem were effectively learned by the convolutional layers of the models and used regardless the differences in the sensors and labelling methods.

Development of a machine learning based classifier includes a large number of elements and processes, hence, all model parameters could not be chosen optimally and some of them were manually fine-tuned or adopted from previous studies. Among these parameters are ones related to the data collection (sampling frequency, number of measured axes, variability of animals and environments), data pre-processing (filter parameters, data fixing and augmentation methods), NN architecture (number, type and size of the layers, number of filters, size of kernels, etc.).

During the experiments, it was found that augmentation simulating sampling loss implemented by Hossain et al. (2020) did not improve performance of the tested models. Because of frequent missing data intervals with length of 10-20 samples, data imputation performed by Weerakody et al. (2021) and Wang Y. et al. (2020) was not effective for this system.

The information missing rate during the data transfer from the tags was about 50%. The main reason is connected with the large number of sent data messages leading to hardware limitations (Tosi et al., 2017). The number of messages depends on the number of tags fitted to cows, and the sampling frequency, which was set to 25Hz. However, the number of cows enrolled in this study was typical for a commercial barn, and this kind of measuring systems should be able to perform in set conditions. Additional analysis of data redundancy must be conducted to find out to what extent the sampling frequency can be decreased in order to diminish the information loss or to increase the number of cows in the same compartment.

In future work, additional uncertainties related to the training of classifiers should be studied. The limits of applicability of trained models should be tested by application on datasets collected in different conditions and environments. To achieve this aim, additional data collection or adoption of existing datasets is required. The influence of the data collected from a specific animal (Wijekoon et al., 2020) on the classification accuracy of the entire herd should be also tested. The minimal amount of data transferred from the sensors should be evaluated by reducing the number of measured acceleration axes and the sampling frequency.

## 5. Conclusion

In this study, we developed a low-cost open-source system with an average F1 score of the feeding behaviour classification equal to 93.9% and analysed different methods and amount of required training data. A dataset of about 20 cow*days for learning using randomly initialized model weights and about 10 cow*hours for the transfer learning was sufficient to achieve F1=90%. Despite of relatively high classification accuracy, additional research is needed to evaluate the applicability of the classifier to other environments and conditions.

## Acknowledgement

The authors acknowledge CSC – IT Center for Science, Finland, for computational resources.

## Funding

This research was funded by ICT-AGRI-2 ERA-NET project “CowBhave: a basic low-cost open-source automated monitoring system for discrimination of dairy cow behavioural activities” funded in Finland by the Ministry of Agriculture and Forestry.

[dataset] Bloch, V., Frondelius, L., Arcidiacono, C., Mancino, M., Pastell, M. 2022. Data from neck acceleration tags for training cow’s feeding behaviour classifier. Zenodo, v1. https://zenodo.org/record/6784671, https://doi.org/10.5281/zenodo.6784671

## Appendix A

To recognise the cow feeding behaviour, different machine learning (ML) methods have been used in research studies, as listed in Table A1. In each study, a specific set of features is used, which was extracted from the acceleration samples in time and frequency domains. However, analysis of the feature types, and amount and physical interpretability have not been found. Studies using NN-based methods are highlighted in bold.

**Table A1.**
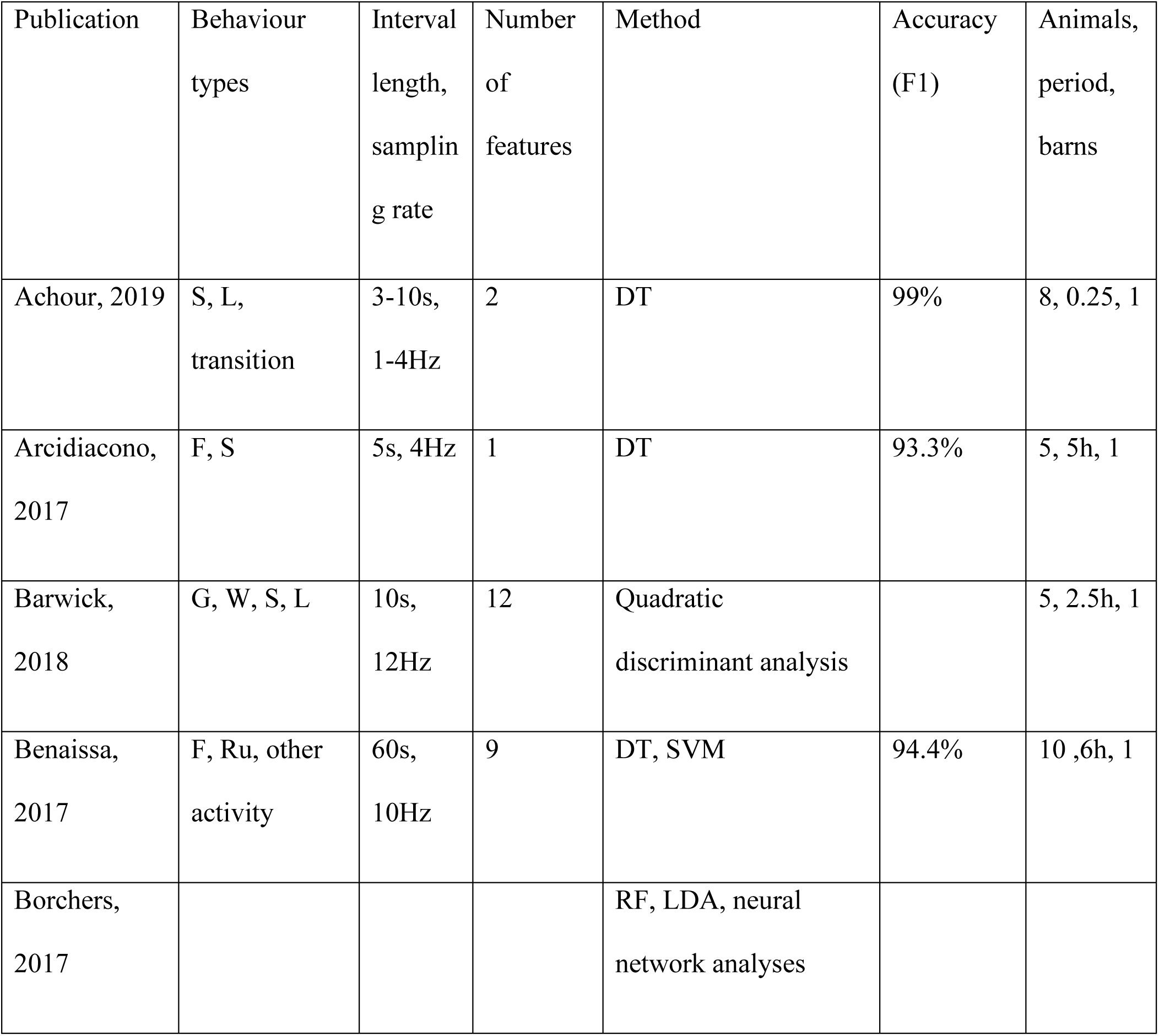
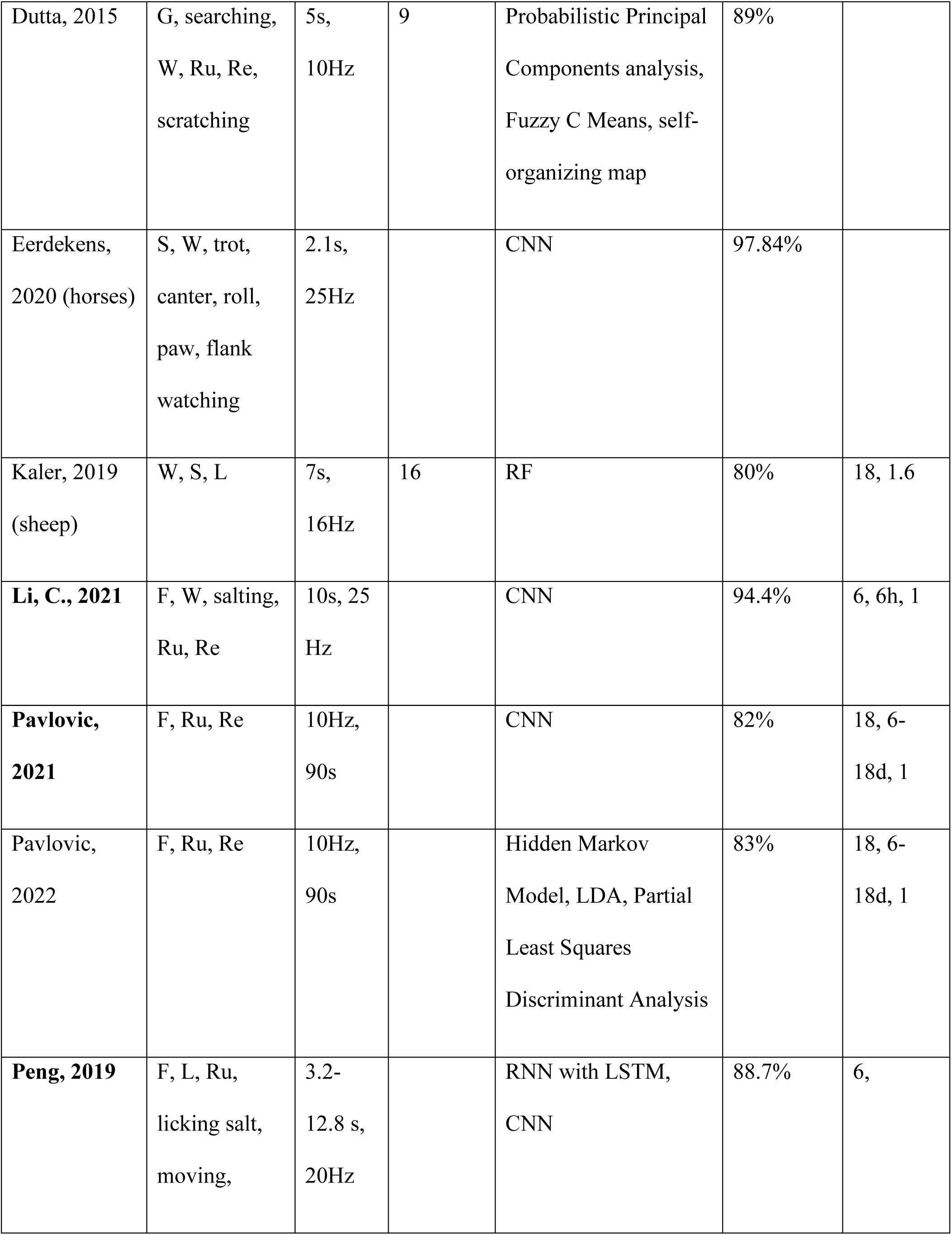
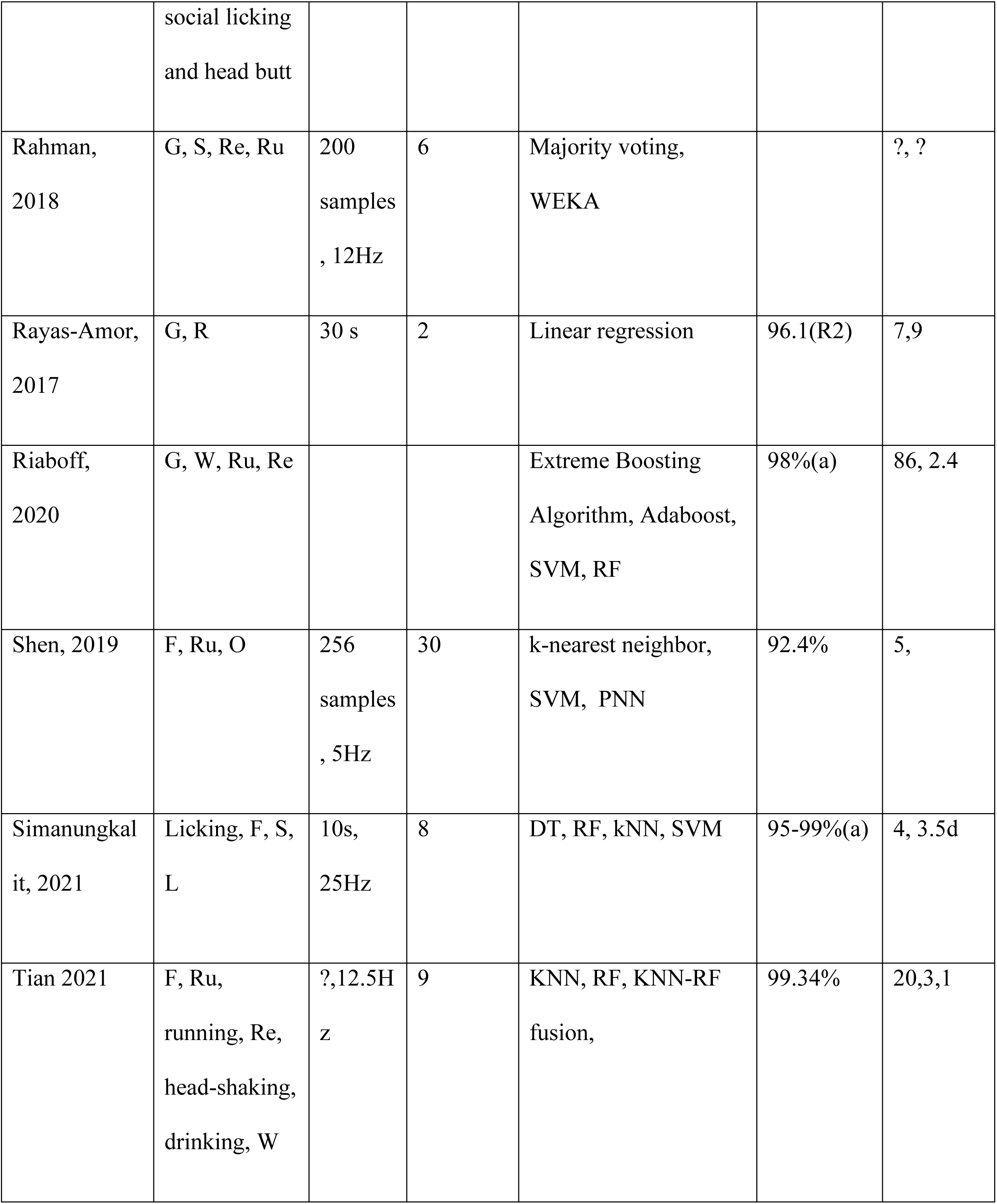
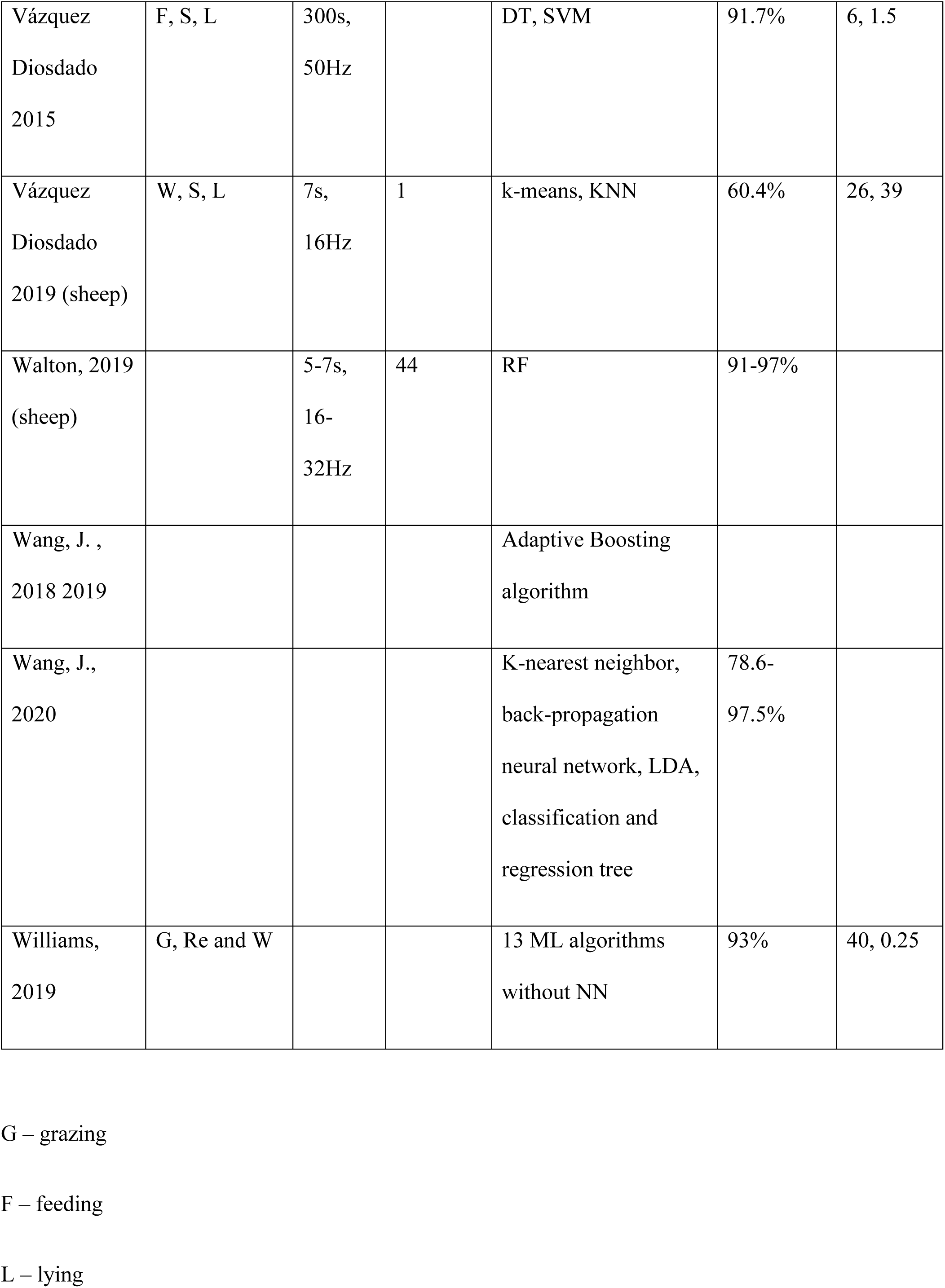
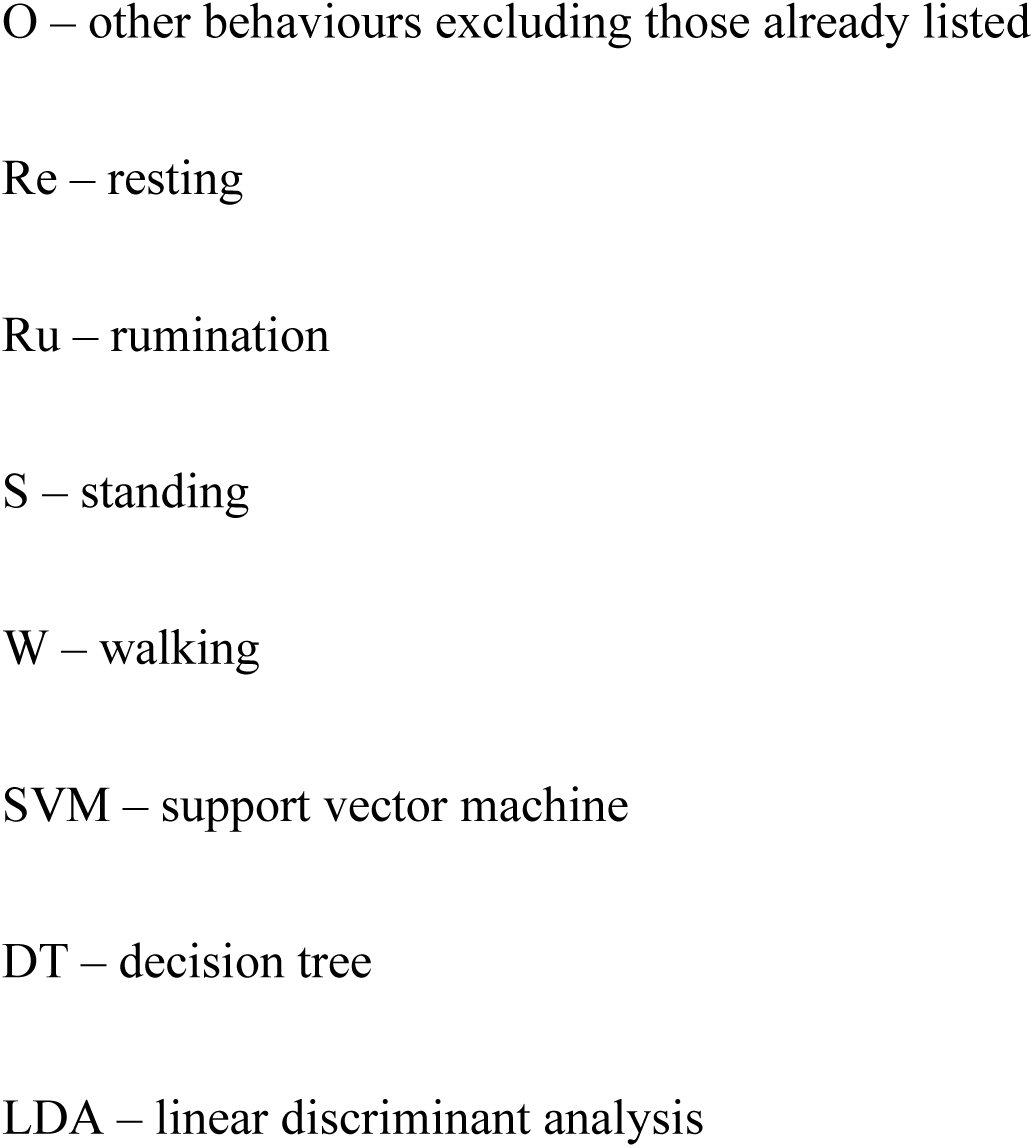
Review of methods for dairy cow behaviour recognition based on acceleration sensors.

## References

1. Achour, B., Belkadi, M., Aoudjit, R., Laghrouche, M. 2019. Unsupervised automated monitoring of dairy cows’ behavior based on Inertial Measurement Unit attached to their back, Computers and Electronics in Agriculture, 167. https://doi.org/10.1016/j.compag.2019.105068

2. Arcidiacono, C., Porto, S.M., Mancino, M., Cascone, G. 2017. Development of a threshold-based classifier for real-time recognition of cow feeding and standing behavioural activities from accelerometer data. Computers and Electronics in Agriculture, 134, 124–134. https://doi.org/10.1016/j.compag.2017.01.021

3. Arcidiacono, C., Mancino, M., Porto, S.M.C., Bloch, V., Pastell, M. 2021. IoT device-based data acquisition system with on-board computation of variables for cow behaviour recognition. Computers and Electronics in Agriculture 191, 106500. https://doi.org/10.1016/j.compag.2021.106500

4. Barwick, J., Lamb, D.W., Dobos, R., Welch, M., Trotter, M. 2018. Categorising sheep activity using a tri-axial accelerometer. Computers and Electronics in Agriculture, 145, 289–297. https://doi.org/10.1016/j.compag.2018.01.007

5. Benaissa, S., Tuyttens, F.A.M., Plets, D., de Pessemier, T., Trogh, J., Tanghe, E., Martens, L., Vandaele, L., van Nuffel, A., Joseph, W., Sonck, B. 2019. On the use of on-cow accelerometers for the classification of behaviours in dairy barns. Research in Veterinary Science, 125, 425–433. https://doi.org/10.1016/j.rvsc.2017.10.005

6. Bloch, V., Pastell, M. 2020. Monitoring of Cow Location in a Barn by an Open-Source, Low-Cost, Low-Energy Bluetooth Tag System. Sensors, 20, 3841. https://doi.org/10.3390/s20143841

7. Borchers, M.R., Chang, Y.M., Proudfoot, K.L., Wadsworth, B.A., Stone, A.E., Bewley, J.M. 2017. Machine-learning-based calving prediction from activity, lying, and ruminating behaviors in dairy cattle. Journal of Dairy Science, 100(7), 5664–5674. https://doi.org/10.3168/jds.2016-11526

8. Brownlee, J. 2016. Master Machine Learning Algorithms: discover how they work and implement them from scratch. Machine Learning Mastery.

9. Dutta, R., Smith, D., Rawnsley, R., Bishop-Hurley, G., Hills, J., Timms, G., Henry, D. 2015. Dynamic cattle behavioural classification using supervised ensemble classifiers. Computers and Electronics in Agriculture, 111, 18–28. https://doi.org/10.1016/j.compag.2014.12.002

10. Eerdekens, A., Deruyck, M., Fontaine, J., Martens, L., De Poorter, E., Joseph, W. 2020. Automatic equine activity detection by convolutional neural networks using accelerometer data. Computers and Electronics in Agriculture, 168. https://doi.org/10.1016/j.compag.2019.105139

11. Eyobu, S.O., Han, D.S. 2018. Feature Representation and Data Augmentation for Human Activity Classification Based on Wearable IMU Sensor Data Using a Deep LSTM Neural Network. Sensors,18(9):2892. https://doi.org/10.3390/s18092892

12. Ferrari, A., Micucci, D., Mobilio, M. Napoletano, P. 2021. Trends in human activity recognition using smartphones. Journal of Reliable Intelligent Environments 7, 189–213. https://doi.org/10.1007/s40860-021-00147-0

13. García, R., Aguilar, J., Toro, M., Pinto, A., Rodríguez, P. 2020. A systematic literature review on the use of machine learning in precision livestock farming. Computers and Electronics in Agriculture, 179. https://doi.org/10.1016/j.compag.2020.105826

14. Grinter, L.N., Campler, M.R., Costa, J.H.C. 2019. Technical note: Validation of a behavior-monitoring collar’s precision and accuracy to measure rumination, feeding, and resting time of lactating dairy cows. Journal of Dairy Science, 102(4), 3487–3494. https://doi.org/10.3168/jds.2018-15563

15. Hossain, T., Ahad, M.A.R., Inoue, S. 2020. A Method for Sensor-Based Activity Recognition in Missing Data Scenario. Sensors 20, 3811. https://doi.org/10.3390/s20143811

16. Kaler, J., Mitsch, J., Vázquez-Diosdado, J.A., Bollard, N., Dottorini, T., Ellis, K.A. 2020. Automated detection of lameness in sheep using machine learning approaches: novel insights into behavioural differences among lame and non-lame sheep. Royal Society Open Science, 7, 190824. https://doi.org/10.1098/rsos.190824

17. Kalouris, G., Zacharaki, E.I., Megalooikonomou, V. 2019. Improving CNN-based activity recognition by data augmentation and transfer learning. IEEE 17th International Conference on Industrial Informatics (INDIN), 1387–1394. https://doi.org/10.1109/INDIN41052.2019.8972135

18. Li, C., Tokgoz, K.K., Fukawa, M., Bartels, J., Ohashi T., Takeda, K., Ito, H. 2021. Data Augmentation for Inertial Sensor Data in CNNs for Cattle Behavior Classification. IEEE Sensors Letters 5(11). https://doi.org/10.1109/LSENS.2021.3119056

19. Li, F., Shirahama, K., Nisar, M.A., Huang, X., Grzegorzek, M. 2020. Deep Transfer Learning for Time Series Data Based on Sensor Modality Classification. Sensors, 20, 4271. https://doi.org/10.3390/s20154271

20. Oh, S., Ashiquzzaman, A., Lee, D., Kim, Y., Kim, J. 2021. Study on Human Activity Recognition Using Semi-Supervised Active Transfer Learning. Sensors, 21, 2760. https://doi.org/10.3390/s21082760

21. Pastell, M., Frondelius, L. 2018. A hidden Markov model to estimate the time dairy cows spend in feeder based on indoor positioning data. Computers and Electronics in Agriculture, 152, 182–185. https://doi.org/10.1016/j.compag.2018.07.005

22. Pavlovic, D., Davison, C., Hamilton, A., Marko, O., Atkinson, R., Michie, C., Crnojević, V., Andonovic, I.; Bellekens, X., Tachtatzis, C. 2021. Classification of Cattle Behaviours Using Neck-Mounted Accelerometer-Equipped Collars and Convolutional Neural Networks. Sensors, 21, 4050. https://doi.org/10.3390/s21124050

23. Pavlovic, D., Czerkawski, M., Davison, C., Marko, O., Michie, C., Atkinson, R., Crnojevic, V., Andonovic, I., Rajovic, V., Kvascev, G., Tachtatzis, C. 2022. Behavioural Classification of Cattle Using Neck-Mounted Accelerometer-Equipped Collars. Sensors, 22(6), 2323. https://doi.org/10.3390/s22062323

24. Peng, Y., Kondo, N., Fujiura, T., Suzuki, T., Wulandari, Yoshioka, H., Itoyama, E. 2019. Classification of multiple cattle behavior patterns using a recurrent neural network with long short-term memory and inertial measurement units. Computers and Electronics in Agriculture, 157, 247–253. https://doi.org/10.1016/j.compag.2018.12.023

25. Rahman, A., Smith, D., Little, B., Ingham, A., Greenwood, P., Bishop-Hurley, G.J. 2018. Cattle behaviour classification from collar, halter, and ear tag sensors. Information Processing in Agriculture, 5(1), 124–133. https://doi.org/10.1016/j.inpa.2017.10.001

26. Rayas-Amor, A.A., Morales-Almaráz, E., Licona-Velázquez, G., Vieyra-Alberto, R., García-Martínez, A., Martínez-García, C.G., Cruz-Monterrosa, R.G., Miranda-de la Lama, G.C. 2017. Triaxial accelerometers for recording grazing and ruminating time in dairy cows: An alternative to visual observations. Journal of Veterinary Behavior, 20,102–108. https://doi.org/10.1016/j.jveb.2017.04.003

27. Riaboff, L., Poggi, S., Madouasse, A., Couvreur, S., Aubin, S., Bédère, N., Goumand, E., Chauvin, A., Plantier, G. 2020. Development of a methodological framework for a robust prediction of the main behaviours of dairy cows using a combination of machine learning algorithms on accelerometer data. Computers and Electronics in Agriculture, 169, 105179. https://doi.org/10.1016/j.compag.2019.105179

28. Riaboff, L. Shalloo, L. Smeaton, A.F. Couvreur, S. Madouasse, A. Keane, M.T. 2022. Predicting livestock behaviour using accelerometers: A systematic review of processing techniques for ruminant behaviour prediction from raw accelerometer data. Computers and Electronics in Agriculture, 192. https://doi.org/10.1016/j.compag.2021.106610

29. Russakovsky, O., Deng, J., Su, H., Krause, J., Satheesh, S., Ma, S., Huang, Z., Karpathy, A., Khosla, A., Bernstein, M., Berg, A. C., Fei-Fei, L. 2015. ImageNet Large Scale Visual Recognition Challenge. International Journal of Computer Vision 115, 211–252. https://doi.org/10.1007/s11263-015-0816-y

30. Shen, W., Cheng, F., Zhang, Y., Wei, X., Fu, Q., Zhang, Y. 2020. Automatic recognition of ingestive-related behaviors of dairy cows based on triaxial acceleration. Information Processing in Agriculture, 7(3), 427–443. https://doi.org/10.1016/j.inpa.2019.10.004

31. Shahriar, M.S., Smith, D.V., Rahman, A., Freeman, M., Hills, J., Rawnsley, R.P., Henry, D., & Bishop-Hurley, G. 2016. Detecting heat events in dairy cows using accelerometers and unsupervised learning. Computers and Electronics in Agriculture, 128, 20–26. https://doi.org/10.1016/j.compag.2016.08.009

32. Simanungkalit, G., Barwick, J., Cowley, F., Dobos, R., Hegarty, R. 2021. A Pilot Study Using Accelerometers to Characterise the Licking Behaviour of Penned Cattle at a Mineral Block Supplement. Animals, 11, 1153. https://doi.org/10.3390/ani11041153

33. Tian, F., Wang, J., Xiong, B., Jiang, L., Song Z., Li, F. 2021. Real-Time Behavioral Recognition in Dairy Cows Based on Geomagnetism and Acceleration Information. IEEE Access, 9, 109497–109509. https://doi.org/10.1109/ACCESS.2021.3099212

34. Tosi, J., Taffoni, F., Santacatterina, M., Sannino, R., Formica, D. 2017. Performance Evaluation of Bluetooth Low Energy: A Systematic Review. Sensors, 17, 2898. https://doi.org/10.3390/s17122898

35. Vázquez Diosdado, J.A., Barker, Z.E., Hodges, H.R., Amory, R.J., Croft, D.P., Bell, N.J., Codling, E.A. 2015. Classification of behaviour in housed dairy cows using an accelerometer-based activity monitoring system. Anima Biotelemetry, 3(1). https://doi.org/10.1186/s40317-015-0045-8

36. Vázquez-Diosdado, J.A., Paul, V., Ellis, K.A., Coates, D., Loomba, R., Kaler, J. A. 2019. Combined Offline and Online Algorithm for Real-Time and Long-Term Classification of Sheep Behaviour: Novel Approach for Precision Livestock Farming. Sensors, 19, 3201. https://doi.org/10.3390/s19143201

37. Um, T.T., Pfister, F.M.J., Pichler, D., Endo, S., Lang, M., Hirche, S., Fietzek, U., Kulić, D. 2017. Data augmentation of wearable sensor data for parkinson’s disease monitoring using convolutional neural networks. In Proceedings of the 19th ACM International Conference on Multimodal Interaction. Association for Computing Machinery, New York, NY, USA, 216–220. https://doi.org/10.1145/3136755.3136817

38. Walton, E., Casey, C., Mitsch, J., Vázquez-Diosdado, J.A., Yan, J., Dottorini, T., Ellis, K.A., Winterlich, A., Kaler, J. 2018 Evaluation of sampling frequency, window size and sensor position for classification of sheep behaviour. Royal Society open science, 5: 171442. http://dx.doi.org/10.1098/rsos.171442

39. Wang, J., Chen, Y., Hao, S., Peng, X., Hu, L. 2019. Deep learning for sensor-based activity recognition: A survey. Pattern Recognition Letters, 119,3–11. https://doi.org/10.1016/j.patrec.2018.02.010

40. Wang, J., He, Z., Zheng, G., Gao, S., Zhao, K. 2018. Development and validation of an ensemble classifier for real-time recognition of cow behavior patterns from accelerometer data and location data. PLOS ONE, 13. https://doi.org/10.1371/journal.pone.0203546

41. Wang, J., Bell, M., Liu, X., Liu, G. 2020. Machine-Learning Techniques Can Enhance Dairy Cow Estrus Detection Using Location and Acceleration Data. Animals, 10(7):1160. https://doi.org/10.3390/ani10071160

42. Wang, Y., Cang, S., Yu, H. 2019. A survey on wearable sensor modality centred human activity recognition in health care. Expert Systems with Applications, 137, 167–190. https://doi.org/10.1016/j.eswa.2019.04.057

43. Weerakody, P.B., Wong, K.W., Wang, G., Ela, W. 2021. A review of irregular time series data handling with gated recurrent neural networks. Neurocomputing, 441, 161–178. https://doi.org/10.1016/j.neucom.2021.02.046

44. Wijekoon, A., Wiratunga, N., Sani, S., Cooper, K. 2020. A knowledge-light approach to personalised and open-ended human activity recognition. Knowledge-Based Systems, 192, 105651. https://doi.org/10.1016/j.knosys.2020.105651

45. Williams, M.L., Wiliam, P.J., Rose, M.T. 2019. Variable segmentation and ensemble classifiers for predicting dairy cow behaviour, Biosystems Engineering, 178, 156–167. https://doi.org/10.1016/j.biosystemseng.2018.11.011

46. WISDM HAR dataset. https://www.cis.fordham.edu/wisdm/dataset.php

