## Supplementary material for "CNN and transfer learning-based classification model for automated cow’s feeding behaviour recognition from accelerometer data": Code: FeedingBehavior_NNlib.py

```
import csv
import numpy
import tensorflow
from tensorflow.keras.models import Sequential
from tensorflow.keras.layers import Dense, Flatten, Dropout, Conv1D, MaxPooling1D, LSTM, TimeDistributed, GlobalAveragePooling1D
from tensorflow.keras.utils import to_categorical
import fnmatch, os
import math
import datetime
from datetime import datetime
from scipy import signal
from scipy import interpolate
from scipy.signal import find_peaks
import matplotlib.pyplot as plt

def CNNModelDefine(ModelName,trainX,trainy):
    n_timesteps, n_features, n_outputs = trainX.shape[1], trainX.shape[2], trainy.shape[1]
    if ModelName=="CNN2": #https://machinelearningmastery.com/cnn-models-for-human-activity-recognition-time-series-classification/
        model = Sequential()
        model.add(Conv1D(filters=64, kernel_size=3+5*0+11*0, activation='relu', input_shape=(n_timesteps,n_features)))
        model.add(Conv1D(filters=64, kernel_size=3+5*0+11*0, activation='relu'))
        model.add(Dropout(0.5))
        model.add(MaxPooling1D(pool_size=2))
        model.add(Flatten())
        model.add(Dense(100, activation='relu'))
        model.add(Dense(n_outputs, activation='softmax'))
        model.compile(loss='categorical_crossentropy', optimizer='adam', metrics=['accuracy'])
    elif ModelName=="LSTMCNN2": #https://machinelearningmastery.com/how-to-develop-rnn-models-for-human-activity-recognition-time-series-classification/
        n_length = 32
        model = Sequential()
        model.add(TimeDistributed(Conv1D(filters=64, kernel_size=3, activation='relu'), input_shape=(None,n_length,n_features)))
        model.add(TimeDistributed(Conv1D(filters=64, kernel_size=3, activation='relu')))
        model.add(TimeDistributed(Dropout(0.5)))
        model.add(TimeDistributed(MaxPooling1D(pool_size=2)))
        model.add(TimeDistributed(Flatten()))
        model.add(LSTM(100))
        model.add(Dropout(0.5))
        model.add(Dense(100, activation='relu'))
        model.add(Dense(n_outputs, activation='softmax'))    
        model.compile(loss='categorical_crossentropy', optimizer='adam', metrics=['accuracy'])
    elif ModelName=="LSTMCNN4": #Ordonez 16, https://doi.org/10.3390/s16010115, https://github.com/STRCWearlab/DeepConvLSTM
        model = Sequential()
        model.add(Conv1D(filters=64, kernel_size=5, activation='relu', input_shape = (n_timesteps, n_features)))
        model.add(Conv1D(filters=64, kernel_size=5, activation='relu'))
        model.add(Conv1D(filters=64, kernel_size=5, activation='relu'))
        model.add(Conv1D(filters=64, kernel_size=5, activation='relu'))
        model.add(Dropout(0.5))
        model.add(LSTM(128, activation='tanh', return_sequences=True))
        model.add(LSTM(128, activation='tanh'))
        model.add(Dense(n_outputs, activation='softmax', dtype='float32'))
        model.compile(loss='categorical_crossentropy', optimizer='adam', metrics=["accuracy", "categorical_accuracy"])
    elif ModelName=="CNN4": #Pavlovic 21, 
        model = Sequential()
        model.add(Conv1D(filters=64, kernel_size=32, strides=1, activation='relu', input_shape = (n_timesteps, n_features)))
        model.add(Conv1D(filters=64, kernel_size=32, strides=2, activation='relu'))
        model.add(Conv1D(filters=64, kernel_size=32, strides=2, activation='relu'))
        model.add(Conv1D(filters=512, kernel_size=1, strides=1, activation='relu'))
        model.add(GlobalAveragePooling1D())
        model.add(Dense(n_outputs, activation='softmax'))
        model.compile(loss='categorical_crossentropy', optimizer='adam', metrics=["accuracy", "categorical_accuracy"])
    else:
        print()
    return model

def ReadLabeledDataFiles(DataFolder,Mask,RefFreq):
    FileList=os.listdir(DataFolder)
    Ax=[]
    Ay=[]
    Az=[]
    Label=[]
    TimeStamp=[]
    CowNo=[]
    TagNo=[]
    for FileName in FileList:
        if FileName.endswith(".csv") and Mask in FileName:
            print(FileName)
            k=FileName.find('_Tag')+4
            m=FileName.find('_Cow')
            l=FileName.find('_',m+2)
            tagno=FileName[k:m]
            cowno=FileName[m+4:l]

            ax=[]
            ay=[]
            az=[]
            label=[]
            timeStamp=[]
            cowNo=[]
            tagNo=[]
            with open(DataFolder+"/"+FileName, mode ='r')as csvfile:
                csvreader = csv.reader(csvfile, delimiter=";")
                rows=[]
                for row in csvreader:
                    rows.append(row)
            
            for i in range(len(rows)):
                cowNo.append(cowno)
                tagNo.append(tagno)
                # TimeStamp.append(rows[i][0])
                timeStamp.append(datetime.strptime(rows[i][0],"%Y-%m-%dT%H:%M:%S.%f").timestamp())
                ax.append(int(rows[i][1]))
                ay.append(int(rows[i][2]))
                az.append(int(rows[i][3]))
                label.append(int(rows[i][4]))
    
            ax, ay, az=AccelerationFiltering(ax, ay, az)
            # ax, ay, az=MissingSampleImputation(ax, ay, az, timeStamp)

            CowNo=list(CowNo)+list(cowNo)
            TagNo=list(TagNo)+list(tagNo)
            TimeStamp=list(TimeStamp)+list(timeStamp)
            Label=list(Label)+list(label)
            Ax=list(Ax)+list(ax)
            Ay=list(Ay)+list(ay)
            Az=list(Az)+list(az)
    
    Label=numpy.asarray(Label)
    CowNo=numpy.asarray(CowNo)
    TagNo=numpy.asarray(TagNo)
    return TagNo, CowNo, TimeStamp, Ax, Ay, Az, Label

def LabeledDataSlicing(Ax, Ay, Az, Label, SliceN, Overlap, TagNo, CowNo, TimeStamp):
    AxSliced=list()
    AySliced=list()
    AzSliced=list()
    LabelSliced=list()

    TagNoSliced=list()
    CowNoSliced=list()
    TimeStampSliced=list()

    N=len(Ax)
    sliceX=numpy.zeros((SliceN, 1))
    sliceY=numpy.zeros((SliceN, 1))
    sliceZ=numpy.zeros((SliceN, 1))
    i=0
    CNprev=0
    while i<N:
        j=0
        L=Label[i]
        CN=CowNo[i]
        while i+j<N and Label[i+j]==L and CowNo[i+j]==CN and j<SliceN:
            sliceX[j]=Ax[i+j]
            sliceY[j]=Ay[i+j]
            sliceZ[j]=Az[i+j]
            j=j+1
        if j==SliceN:
            sliceX=AccNormalization(sliceX)
            sliceY=AccNormalization(sliceY)
            sliceZ=AccNormalization(sliceZ)
            AxSliced.append(sliceX.copy())
            AySliced.append(sliceY.copy())
            AzSliced.append(sliceZ.copy())
            LabelSliced.append(L)
            CowNoSliced.append(CN)
            TagNoSliced.append(TagNo[i])
            TimeStampSliced.append(TimeStamp[i])
        i=i+int(SliceN*Overlap)

    return AxSliced, AySliced, AzSliced, LabelSliced, TagNoSliced, CowNoSliced, TimeStampSliced

def AccelerationFiltering(Ax, Ay, Az):#WindowN,
    Draw=False#True#
    if Draw:
        plt.plot(range(len(Ax)),Ax, 'r*')
        plt.plot(range(len(Ax)),Ay, 'g*')
        plt.plot(range(len(Ax)),Az, 'b*')

    q=[]
    AX=[]
    AY=[]
    AZ=[]
    for i in range(len(Ax)):
        if Ax[i]!=0:
            AX.append(Ax[i])
            AY.append(Ay[i])
            AZ.append(Az[i])
            q.append(i)
    
    b = signal.firwin(511, cutoff = 0.1, fs = 25, window = "hamming", pass_zero="highpass")
    AX = signal.lfilter(b, 1.0, AX)
    AY = signal.lfilter(b, 1.0, AY)
    AZ = signal.lfilter(b, 1.0, AZ)

    for i in range(len(q)):
        Ax[q[i]]=AX[i]
        Ay[q[i]]=AY[i]
        Az[q[i]]=AZ[i]
    
    if Draw:
        plt.plot(range(len(Ax)),Ax,'b.')
        plt.plot(range(len(Ax)),Ay,'r.')
        plt.plot(range(len(Ax)),Az,'g.')
        plt.show()

    return Ax, Ay, Az

def MissingSampleImputation(Ax, Ay, Az, TimeStamp):#, Label, TagNo, CowNo
    Draw=False#True#
    if Draw:
        plt.plot(TimeStamp,Ax, 'r*')
        plt.plot(TimeStamp,Ay, 'g*')
        plt.plot(TimeStamp,Az, 'b*')

    q=[]
    AX=[]
    AY=[]
    AZ=[]
    TS=[]
    # q.append(0)
    # AX.append(Ax[0])
    # AY.append(Ay[0])
    # AZ.append(Az[0])
    # TS.append(TimeStamp[0])
    for i in range(1,len(Ax)-1):
        if Ax[i]!=0:
            AX.append(Ax[i])
            AY.append(Ay[i])
            AZ.append(Az[i])
            TS.append(TimeStamp[i])
            q.append(i)

    f=interpolate.interp1d(TS,AX,kind='cubic',fill_value="extrapolate")
    Ax=f(TimeStamp)
    f=interpolate.interp1d(TS,AY,kind='cubic',fill_value="extrapolate")
    Ay=f(TimeStamp)
    f=interpolate.interp1d(TS,AZ,kind='cubic',fill_value="extrapolate")
    Az=f(TimeStamp)
    # # f=interpolate.interp1d(TimeStamp,TagNo,kind='nearest')
    # # TagNo=f(TimeStampNew)
    # # f=interpolate.interp1d(TimeStamp,CowNo,kind='nearest')
    # # CowNo=f(TimeStampNew)
    # # f=interpolate.interp1d(TimeStamp,Label,kind='nearest')
    # # Label=f(TimeStampNew)

    if Draw:
        plt.plot(TimeStamp,Ax,'g.')
        plt.plot(TimeStamp,Ay,'b.')
        plt.plot(TimeStamp,Az,'r.')

        # # plt.figure()
        # # plt.plot(TimeStamp,TagNo, 'k.')
        # # plt.plot(TimeStamp,CowNo, 'k*')

        plt.show()

    return Ax, Ay, Az

def AccNormalization(A):
    # return A/numpy.std(A)
    return A/1024

def DataBalance(Ax, Ay, Az, Label, CowNo):#TimeStamp
    N=len(Label)
    m=max(Label)

    (unique, LabelCounts) = numpy.unique(Label, return_counts=True)
    print("Total class distribution:"+str(LabelCounts))
    LabelN=len(LabelCounts)

    CowNoList=list(set(CowNo))
    CowNo=numpy.array(CowNo)
    Ax=numpy.array(Ax)
    Ay=numpy.array(Ay)
    Az=numpy.array(Az)
    Label=numpy.array(Label)
    # TimeStamp=numpy.array(TimeStamp)
    AxBalanced=list()
    AyBalanced=list()
    AzBalanced=list()
    LabelBalanced=list()
    CowNoBalanced=list()

    for Cow_i in range(len(CowNoList)):
        q=numpy.argwhere(CowNo==CowNoList[Cow_i])
        AxCowNo_i=Ax[q]
        AyCowNo_i=Ay[q]
        AzCowNo_i=Az[q]
        LabelCowNo_i=Label[q]

        for i in range(len(LabelCowNo_i)):
            AxBalanced.append(numpy.squeeze(AxCowNo_i[i]))
            AyBalanced.append(numpy.squeeze(AyCowNo_i[i]))
            AzBalanced.append(numpy.squeeze(AzCowNo_i[i]))
            LabelBalanced.append(LabelCowNo_i[i][0])
            CowNoBalanced.append(CowNoList[Cow_i])

        (unique, LabelCounts) = numpy.unique(LabelCowNo_i, return_counts=True)
        print("Cow "+str(CowNoList[Cow_i])+", classes "+str(LabelCounts))
        LabelCountsMax=numpy.max(LabelCounts)
        for Label_i in range(LabelN):
            e=numpy.argwhere(LabelCowNo_i==(Label_i+1))
            AxCowNo_iLabel_i=AxCowNo_i[e[:,0]]
            AyCowNo_iLabel_i=AyCowNo_i[e[:,0]]
            AzCowNo_iLabel_i=AzCowNo_i[e[:,0]]
            for j in range(LabelCountsMax-LabelCounts[Label_i]):
                r=numpy.random.randint(LabelCounts[Label_i])
                ax0=numpy.squeeze(AxCowNo_iLabel_i[r])
                ay0=numpy.squeeze(AyCowNo_iLabel_i[r])
                az0=numpy.squeeze(AzCowNo_iLabel_i[r])
                RotA=math.pi*numpy.random.rand()
                s=math.sin(RotA)
                c=math.cos(RotA)
                ax=ax0
                ay=ay0*c-az0*s
                az=ay0*s+az0*c
                AxBalanced.append(ax)
                AyBalanced.append(ay)
                AzBalanced.append(az)
                LabelBalanced.append(Label_i+1)
                CowNoBalanced.append(CowNoList[Cow_i])

    (unique, LabelCounts) = numpy.unique(LabelBalanced, return_counts=True)
    print("Balanced class distribution: "+str(LabelCounts))

    return AxBalanced, AyBalanced, AzBalanced, LabelBalanced, CowNoBalanced

def DataSlicing(Ax, Ay, Az, Timestamp, n):
    AxSliced=list()
    AySliced=list()
    AzSliced=list()
    TimestampSliced=list()
    N=len(Ax)
    sliceX=numpy.zeros((n, 1))
    sliceY=numpy.zeros((n, 1))
    sliceZ=numpy.zeros((n, 1))
    i=0
    while i+n<N:
        for j in range(n):
            sliceX[j]=Ax[i+j]
            sliceY[j]=Ay[i+j]
            sliceZ[j]=Az[i+j]
        sliceX=AccNormalization(sliceX)
        sliceY=AccNormalization(sliceY)
        sliceZ=AccNormalization(sliceZ)
        AxSliced.append(sliceX.copy())
        AySliced.append(sliceY.copy())
        AzSliced.append(sliceZ.copy())
        TimestampSliced.append(Timestamp[i])#+n//2
        i=i+j

    return AxSliced, AySliced, AzSliced, TimestampSliced

def AccRotationAugmentation(Ax0, Ay0, Az0, Label0, CowNo0):
    Ax=list()
    Ay=list()
    Az=list()
    Label=list()
    CowNo=list()

    for i in range(len(Ax0)):
        ax0=Ax0[i]
        ay0=Ay0[i]
        az0=Az0[i]

        RotA=2*math.pi*numpy.random.rand()
        s=math.sin(RotA)
        c=math.cos(RotA)

        ax=ax0
        ay=ay0*c-az0*s
        az=ay0*s+az0*c
        Ax.append(ax)
        Ay.append(ay)
        Az.append(az)
        Label.append(Label0[i])
        CowNo.append(CowNo0[i])

    (unique, LabelCounts) = numpy.unique(Label, return_counts=True)
    # print("Augmented rotation class distribution: "+str(LabelCounts))

    return Ax, Ay, Az, Label, CowNo

def AccSamplingAugmentation(Ax, Ay, Az, Label, CowNo, MaxMissingSamplingRate):
    n=numpy.shape(Ax)
    WindowSize=n[1]
    k=0.05
    MaxMissingSamples=int(MaxMissingSamplingRate*WindowSize)
    MinMissingSamples=int(0.05*WindowSize)

    for i in range(len(Ax)):
        ax0=Ax[i]
        ay0=Ay[i]
        az0=Az[i]

        # plt.plot(range(WindowSize), ax0,'b*')

        d=numpy.count_nonzero(ax0==0)
        if d<WindowSize*k:
            MissingRate=int(numpy.random.randint(MinMissingSamples,MaxMissingSamples))
            print(MissingRate)
            for j in range(int(MissingRate/5)):
                w=numpy.random.randint(0, WindowSize-1-5)
                # print(w)
                ax0[w]=0
                ay0[w]=0
                az0[w]=0
                ax0[w+1],ax0[w+2],ax0[w+3],ax0[w+4]=0,0,0,0
                ay0[w+1],ay0[w+2],ay0[w+3],ay0[w+4]=0,0,0,0
                az0[w+1],az0[w+2],az0[w+3],az0[w+4]=0,0,0,0
            Ax.append(ax0)
            Ay.append(ay0)
            Az.append(az0)
            Label.append(Label[i])
            CowNo.append(CowNo[i])
        # plt.plot(range(WindowSize), ax0,'r.')
        # plt.show()

    (unique, LabelCounts) = numpy.unique(Label, return_counts=True)
    print("Augmented sampling class distribution: "+str(LabelCounts))

    return Ax, Ay, Az, Label, CowNo

from sklearn import metrics
def PresentPerformance(LabelReference,LabelPredicted,Classes,ClassNames):
    N=len(LabelPredicted)
    ClassN=len(Classes)
    Ttotal = 0
    TP=numpy.zeros((ClassN, 1))
    TN=numpy.zeros((ClassN, 1))
    FP=numpy.zeros((ClassN, 1))
    FN=numpy.zeros((ClassN, 1))
    for i in range(N):
        if LabelPredicted[i]==LabelReference[i]:
            Ttotal=Ttotal+1

        for j in range(ClassN):
            if LabelReference[i]==Classes[j]:
                if LabelPredicted[i]==Classes[j]:
                    TP[j]=TP[j]+1
                else:
                    FN[j]=FN[j]+1
            else:
                if LabelPredicted[i]==Classes[j]:
                    FP[j]=FP[j]+1
                else:
                    TN[j]=TN[j]+1

    PerfVect=numpy.zeros((ClassN, 3))
    print()
    TotAcc=round(Ttotal/N*100,1)
    print('Total accuracy =',TotAcc,'%, N =',N)
    for j in range(ClassN):
        sens=numpy.squeeze(TP[j]/(TP[j]+FP[j]))
        spec=numpy.squeeze(TN[j]/(TN[j]+FP[j]))
        F1=2*sens*spec/(sens+spec)
        print(ClassNames[j],' sens =',numpy.round(sens*100,2),'%, spec =',numpy.round(spec*100,2),'%, F1 =',numpy.round(F1*100,2),'%, N =',numpy.squeeze(int(TP[j]+TN[j])))
        PerfVect[j]=[sens, spec, F1]

    ConfusionMatr=numpy.zeros((3, 3))
    for i in range(N):
        ConfusionMatr[int(LabelPredicted[i])-1,int(LabelReference[i])-1]+=1

    print()
    print(numpy.squeeze(ClassNames))
    # print(numpy.squeeze(ConfusionMatr.astype(int)))

    print(metrics.confusion_matrix(LabelReference,LabelPredicted))
    print(metrics.classification_report(LabelReference,LabelPredicted, digits=3))

    return PerfVect, ConfusionMatr, TotAcc

from scipy.stats import skew, kurtosis
def AccelerationFeatures(Ax,Ay,Az,Label,fs):
    # Ax_Mean=numpy.mean(Ax, axis=1)
    # Ay_Mean=numpy.mean(Ay, axis=1)
    # Az_Mean=numpy.mean(Az, axis=1)
    # Ax_STD=numpy.std(Ax, axis=1)
    # Ay_STD=numpy.std(Ay, axis=1)
    # Az_STD=numpy.std(Az, axis=1)
    Ax_Mean=numpy.mean(Ax)
    Ay_Mean=numpy.mean(Ay)
    Az_Mean=numpy.mean(Az)
    Ax_STD=numpy.std(Ax)
    Ay_STD=numpy.std(Ay)
    Az_STD=numpy.std(Az)
    Ax_Range=max(Ax)-min(Ax)
    Ay_Range=max(Ay)-min(Ay)
    Az_Range=max(Az)-min(Az)
    Ax_Skew=skew(Ax)
    Ay_Skew=skew(Ay)
    Az_Skew=skew(Az)
    Ax_Kurt=kurtosis(Ax)
    Ay_Kurt=kurtosis(Ay)
    Az_Kurt=kurtosis(Az)
    Ax_Zero_crossings = len(numpy.where(numpy.diff(numpy.signbit(Ax-Ax_Mean)))[0])
    Ay_Zero_crossings = len(numpy.where(numpy.diff(numpy.signbit(Ay-Ay_Mean)))[0])
    Az_Zero_crossings = len(numpy.where(numpy.diff(numpy.signbit(Az-Az_Mean)))[0])
    Ax_Signal_area=sum(Ax)/fs
    Ay_Signal_area=sum(Ay)/fs
    Az_Signal_area=sum(Az)/fs
    f, Ax_Pxx_den = signal.periodogram(Ax, fs)
    f, Ay_Pxx_den = signal.periodogram(Ay, fs)
    f, Az_Pxx_den = signal.periodogram(Az, fs)

    b, a = signal.butter(5, 25, 'low', fs=100)
    Ax_Pxx_den_f = signal.filtfilt(b, a, Ax_Pxx_den)
    Ay_Pxx_den_f = signal.filtfilt(b, a, Ay_Pxx_den)
    Az_Pxx_den_f = signal.filtfilt(b, a, Az_Pxx_den)

    # Ax_Spectral_entropy=sum(Ax_Pxx_den*numpy.log(Ax_Pxx_den_f))
    # Ay_Spectral_entropy=sum(Ay_Pxx_den*numpy.log(Ay_Pxx_den_f))
    # Az_Spectral_entropy=sum(Az_Pxx_den*numpy.log(Az_Pxx_den_f))

    # Ax_Dominant_frequency=numpy.argmax(Ax_Pxx_den_f)
    # Ay_Dominant_frequency=numpy.argmax(Ay_Pxx_den_f)
    # Az_Dominant_frequency=numpy.argmax(Az_Pxx_den_f)

    Ax_Spectral_area=sum(Ax_Pxx_den_f)
    Ay_Spectral_area=sum(Ay_Pxx_den_f)
    Az_Spectral_area=sum(Az_Pxx_den_f)

    Ax_Maxima, _ = find_peaks(Ax_Pxx_den_f, distance=30)
    sort_index = numpy.argsort(-Ax_Pxx_den_f[Ax_Maxima])
    if len(sort_index)>0:
        Ax_Dominant_frequency=Ax_Maxima[sort_index[0]]
    else:
        Ax_Dominant_frequency=0
    if len(sort_index)>1:
        Ax_2_frequency=Ax_Maxima[sort_index[1]]
    else:
        Ax_2_frequency=0
    if len(sort_index)>2:
        Ax_3_frequency=Ax_Maxima[sort_index[2]]
    else:
        Ax_3_frequency=0

    Ay_Maxima, _ = find_peaks(Ay_Pxx_den_f, distance=30)
    sort_index = numpy.argsort(-Ay_Pxx_den_f[Ay_Maxima])
    if len(sort_index)>0:
        Ay_Dominant_frequency=Ay_Maxima[sort_index[0]]
    else:
        Ay_Dominant_frequency=0
    if len(sort_index)>1:
        Ay_2_frequency=Ay_Maxima[sort_index[1]]
    else:
        Ay_2_frequency=0
    if len(sort_index)>2:
        Ay_3_frequency=Ay_Maxima[sort_index[2]]
    else:
        Ay_3_frequency=0

    Az_Maxima, _ = find_peaks(Az_Pxx_den_f, distance=30)
    sort_index = numpy.argsort(-Az_Pxx_den_f[Az_Maxima])
    if len(sort_index)>0:
        Az_Dominant_frequency=Az_Maxima[sort_index[0]]
    else:
        Az_Dominant_frequency=0
    if len(sort_index)>1:
        Az_2_frequency=Az_Maxima[sort_index[1]]
    else:
        Az_2_frequency=0
    if len(sort_index)>2:
        Az_3_frequency=Az_Maxima[sort_index[2]]
    else:
        Az_3_frequency=0
    
    Features=numpy.dstack((Ax_Mean,Ay_Mean,Az_Mean))
    Features=numpy.dstack((Features,Ax_STD,Ay_STD,Az_STD))
    Features=numpy.dstack((Features,Ax_Range,Ay_Range,Az_Range))
    Features=numpy.dstack((Features,Ax_Skew,Ay_Skew,Az_Skew))
    Features=numpy.dstack((Features,Ax_Kurt,Ay_Kurt,Az_Kurt))
    Features=numpy.dstack((Features,Ax_Zero_crossings,Ay_Zero_crossings,Az_Zero_crossings))
    Features=numpy.dstack((Features,Ax_Signal_area,Ay_Signal_area,Az_Signal_area))
    # Features=numpy.dstack((Features,Ax_Spectral_entropy,Ay_Spectral_entropy,Az_Spectral_entropy))
    Features=numpy.dstack((Features,Ax_Dominant_frequency,Ay_Dominant_frequency,Az_Dominant_frequency))
    Features=numpy.dstack((Features,Ax_Spectral_area,Ay_Spectral_area,Az_Spectral_area))
    Features=numpy.dstack((Features,Ax_2_frequency,Ay_2_frequency,Az_2_frequency))
    Features=numpy.dstack((Features,Ax_3_frequency,Ay_3_frequency,Az_3_frequency))

    # plt.plot(Ax,'r*')
    # plt.figure()
    # plt.plot(Ax_Pxx_den, 'k.')
    # plt.plot(Ax_Pxx_den_f, 'g')
    # print(Ax_Maxima)
    # plt.plot(Ax_Maxima,Ax_Pxx_den_f[Ax_Maxima],'b*')
    # plt.plot(Ax_Dominant_frequency,Ax_Pxx_den_f[Ax_Dominant_frequency],'ro')
    # plt.plot(Ax_2_frequency,Ax_Pxx_den_f[Ax_2_frequency],'go')
    # plt.plot(Ax_3_frequency,Ax_Pxx_den_f[Ax_3_frequency],'bo')
    # plt.show()
    # print(Features)
    # print(Ax_Zero_crossings)

    Features=numpy.squeeze(Features)
    Label=numpy.squeeze(Label)
    return Features,Label
```
