## Supplementary material for "CNN and transfer learning-based classification model for automated cow’s feeding behaviour recognition from accelerometer data": Code: Main_FeedingBehaviour_CNNModelBuilding.py

```
# .\venv\Scripts\activate
# cd C:\Users\03138529\Dropbox\Luke\CowBhave
# python D:/CowBhave1/Main_FeedingBehaviour_CNNModelBuilding.py

from FeedingBehavior_NNlib import *
import fnmatch, os
from sklearn.model_selection import train_test_split
import numpy
import matplotlib.pyplot as plt
import joblib
from datetime import datetime
import scipy.signal
import h5py
import sys

Freq=25 #Hz
FeedingM, RuminatingM, NothingM, DrinkingM = 1, 2, 3, 4
ProjectFolder="D:/CowBhave1/"
ModelFolder=ProjectFolder+"Models25"
if not os.path.exists(ModelFolder):
    os.mkdir(ModelFolder)

ModelType="CNN2"
print(ModelType)
DataSetFolder,DataSetName=[],[]
# DataSetFolder.append(ProjectFolder+"Labeled25"),DataSetName.append("Barn2")
DataSetFolder.append(ProjectFolder+"Pavlovic21/Labeled25"),DataSetName.append("BarnP")
# DataSetFolder.append(ProjectFolder+"Labeled25_60B1"),DataSetName.append("Barn2_60B1")
# DataSetFolder.append(ProjectFolder+"Labeled25_60B2"),DataSetName.append("Barn2_60B2")
# DataSetFolder.append(ProjectFolder+"Labeled25_60B3"),DataSetName.append("Barn2_60B3")
# DataSetFolder.append(ProjectFolder+"Labeled25_60B4"),DataSetName.append("Barn2_60B4")
# DataSetFolder.append(ProjectFolder+"Labeled25_60B5"),DataSetName.append("Barn2_60B5")
# DataSetFolder.append(ProjectFolder+"Labeled25_60B6"),DataSetName.append("Barn2_60B6")
# DataSetFolder.append(ProjectFolder+"Labeled25_60B7"),DataSetName.append("Barn2_60B7")
# DataSetFolder.append(ProjectFolder+"Labeled25_60B8"),DataSetName.append("Barn2_60B8")
# DataSetFolder.append(ProjectFolder+"Labeled25_60B9"),DataSetName.append("Barn2_60B9")
# DataSetFolder.append(ProjectFolder+"Labeled25_60B10"),DataSetName.append("Barn2_60B10")
# DataSetFolder.append(ProjectFolder+"Labeled25_5"),DataSetName.append("Barn2_5")
# DataSetFolder.append(ProjectFolder+"Labeled25_10"),DataSetName.append("Barn2_10")
# DataSetFolder.append(ProjectFolder+"Labeled25_20"),DataSetName.append("Barn2_20")
# DataSetFolder.append(ProjectFolder+"Labeled25_30"),DataSetName.append("Barn2_30")
# DataSetFolder.append(ProjectFolder+"Labeled25_40"),DataSetName.append("Barn2_40")
# DataSetFolder.append(ProjectFolder+"Labeled25_50"),DataSetName.append("Barn2_50")
# DataSetFolder.append(ProjectFolder+"Labeled25_60"),DataSetName.append("Barn2_60")
# DataSetFolder.append(ProjectFolder+"Labeled25_70"),DataSetName.append("Barn2_70")
# DataSetFolder.append(ProjectFolder+"Labeled25_80"),DataSetName.append("Barn2_80")
# DataSetFolder.append(ProjectFolder+"Labeled25_90"),DataSetName.append("Barn2_90")
WindowSizeList=[5,10,20,30,60,90,120,180,300] #sec
# WindowSizeList=[60] #sec
FoldN=2

Vers_i=0

for DataSetFolder_i in range(len(DataSetFolder)):
    for WindowSize in WindowSizeList:
        for Fold_i in range(FoldN+1):
            if Fold_i==0:
                DataFileName=DataSetFolder[DataSetFolder_i]+"/"+'FeedingBehaviour_Training'+"_WS"+str(WindowSize)+"_F"+str(Freq)+'.h5'
                ModelFileName=ModelFolder+"/"+ModelType+DataSetName[DataSetFolder_i]+"WS"+str(WindowSize)+"Fold"+str(0)+"V"+str(Vers_i)
            else:
                DataFileName=DataSetFolder[DataSetFolder_i]+"/"+'FeedingBehaviour_Training'+"_WS"+str(WindowSize)+"_F"+str(Freq)+"_Fold"+str(Fold_i)+'.h5'
                ModelFileName=ModelFolder+"/"+ModelType+DataSetName[DataSetFolder_i]+"WS"+str(WindowSize)+"Fold"+str(Fold_i)+"V"+str(Vers_i)

            print(DataFileName)
            f = h5py.File(DataFileName, 'r')
            ASlicedTraining = f['AccXYZ'][...]
            LabelSlicedOneHotTraining = f['LabelOneHot'][...]
            LabelSlicedTraining = f['Label'][...]
            f.close()

            print(len(LabelSlicedTraining))

            print(ModelFileName)
            epochs, batch_size, verbose = 3+50*0, 32, 1
            model=CNNModelDefine(ModelType,ASlicedTraining,LabelSlicedOneHotTraining)
            print(model.summary())
            train_history = model.fit(ASlicedTraining, LabelSlicedOneHotTraining, epochs=epochs, batch_size=batch_size, verbose=verbose)
            model.save(ModelFileName)

            print(train_history.history['loss'])
            p=train_history.history['loss']
            with open(ModelFileName+'/train_history.csv', 'w') as f:
                for item in p:
                    f.write("%s\n" % item)
print('\a')
```
