## Supplementary material for "CNN and transfer learning-based classification model for automated cow’s feeding behaviour recognition from accelerometer data": Code: Main_FeedingBehaviour_CNNSelfValidation.py

```
# .\venv\Scripts\activate
# cd C:\Users\03138529\Dropbox\Luke\CowBhave
# python Main_FeedingBehaviour_CNNSelfValidation.py

from FeedingBehavior_NNlib import *
import fnmatch, os
from sklearn.model_selection import train_test_split
from sklearn.ensemble import RandomForestClassifier
from sklearn import svm
import numpy
import matplotlib.pyplot as plt
import joblib
from datetime import datetime
import scipy.signal
import h5py

# ProjectFolder="/scratch/project_2001335/CowBhave/"
ProjectFolder="D:/CowBhave1/"
ModelFolder=ProjectFolder+"Models25"
WindowSizeList=[5,10,20,30,60,90,120,180,300] #sec
# WindowSizeList=[60] #sec
Freq=25 #Hz
FeedingM, RuminatingM, NothingM, DrinkingM = 1, 2, 3, 4

DataSetFolder,DataSetName=[],[]
# DataSetFolder.append(ProjectFolder+"Data_Exp08_11_2019/Labeled10"),DataSetName.append("Barn1")
DataSetFolder.append(ProjectFolder+"Data_Exp15_02_2021/Labeled25"),DataSetName.append("Barn2")
# DataSetFolder.append(ProjectFolder+"Pavlovic21/Labeled25"),DataSetName.append("BarnP")
FoldN=10
ModelType="CNN2"
PretrainedModelName="BarnP"
FrozenLayersN=2

Vers_i=0
for WindowSize in WindowSizeList:
    for DataSetFolder_i in range(len(DataSetFolder)):
        for Fold_i in range(FoldN+1):
            if Fold_i==0:
                DataFileName=DataSetFolder[DataSetFolder_i]+"/"+'FeedingBehaviour_Validation'+"_WS"+str(WindowSize)+"_F"+str(Freq)+'.h5'
                # ModelFileName=ModelFolder+"/"+ModelType+DataSetName[DataSetFolder_i]+"WS"+str(WindowSize)+"Fold"+str(0)+"V"+str(Vers_i)
                ModelFileName=ModelFolder+"/"+ModelType+PretrainedModelName+"WS"+str(WindowSize)+"Fold"+str(0)+"V"+str(Vers_i)+"_TL"+DataSetName[DataSetFolder_i]+"_L"+str(FrozenLayersN)+"Fold"+str(0)+"V"+str(Vers_i)
            else:
                DataFileName=DataSetFolder[DataSetFolder_i]+"/"+'FeedingBehaviour_Validation'+"_WS"+str(WindowSize)+"_F"+str(Freq)+"_Fold"+str(Fold_i)+'.h5'
                # ModelFileName=ModelFolder+"/"+ModelType+DataSetName[DataSetFolder_i]+"WS"+str(WindowSize)+"Fold"+str(Fold_i)+"V"+str(Vers_i)
                ModelFileName=ModelFolder+"/"+ModelType+PretrainedModelName+"WS"+str(WindowSize)+"Fold"+str(0)+"V"+str(Vers_i)+"_TL"+DataSetName[DataSetFolder_i]+"_L"+str(FrozenLayersN)+"Fold"+str(Fold_i)+"V"+str(Vers_i)

            print(DataFileName)
            f = h5py.File(DataFileName, 'r')
            ASlicedValidation = f['AccXYZ'][...]
            LabelSlicedValidation = f['Label'][...]
            f.close()

            print(ModelFileName)
            model=tensorflow.keras.models.load_model(ModelFileName)

            FeedingBehaviorPredictedP=model.predict(ASlicedValidation)
            FeedingBehaviorPredicted=list()
            for i in range(len(FeedingBehaviorPredictedP)):
                m,k=MaxInd(FeedingBehaviorPredictedP[i])
                FeedingBehaviorPredicted.append(k+1)

            FeedingBehaviorPredicted=scipy.signal.medfilt(FeedingBehaviorPredicted,5)
            PerfVect, ConfusionMatr, TotAcc=PresentPerformance(LabelSlicedValidation,FeedingBehaviorPredicted,[FeedingM, RuminatingM, NothingM],["Feeding", "Ruminating", "Nothing"])
            now = datetime.now()
            dt_string = now.strftime("%Y%m%d_%H%M%S")
            # s=dt_string+";"+ModelType+";"+DataSetName[DataSetFolder_i]+";"+DataSetName[DataSetFolder_i]+";"+str(WindowSize)+";"+str(Fold_i)+";"+str(Vers_i)+";"+str(TotAcc)+";"
            s=dt_string+";"+ModelType+";"+PretrainedModelName+"L"+str(FrozenLayersN)+"TL"+DataSetName[DataSetFolder_i]+";"+DataSetName[DataSetFolder_i]+";"+str(WindowSize)+";"+str(Fold_i)+";"+str(Vers_i)+";"+str(TotAcc)+";"
            for i in range(len(ConfusionMatr)):
                for j in range(len(ConfusionMatr)):
                    s=s+str(ConfusionMatr[i,j])+";"
            fileH=open(ModelFolder+"/"+'Performance.csv', 'a')
            fileH.write(s+'\n')
            fileH.close()

print('\a')
```
