## Supplementary material for "CNN and transfer learning-based classification model for automated cow’s feeding behaviour recognition from accelerometer data": Code: Main_FeedingBehaviour_CNNTransferLearning.py

```
# .\venv\Scripts\activate
# cd C:\Users\03138529\Dropbox\Luke\CowBhave
# python D:/CowBhave1/Main_FeedingBehaviour_CNNTransferLearning.py
# https://machinelearningmastery.com/how-to-improve-performance-with-transfer-learning-for-deep-learning-neural-networks/

from FeedingBehavior_NNlib import *
import fnmatch, os
from sklearn.model_selection import train_test_split
from sklearn.ensemble import RandomForestClassifier
from sklearn import svm
import numpy
import matplotlib.pyplot as plt
import joblib
from datetime import datetime
import scipy.signal
import h5py
import sys

Freq=25 #Hz
FeedingM, RuminatingM, NothingM, DrinkingM = 1, 2, 3, 4
# ProjectFolder="/scratch/project_2001335/CowBhave/"
ProjectFolder="D:/CowBhave1/"
ModelFolder=ProjectFolder+"Models25"

ModelType="CNN2"
FrozenLayersN=1
WindowSizeList=[5,10,20,30,60,90,120,180,300] #sec
# WindowSizeList=[60] #sec
DataSetFolder,DataSetName=[],[]
DataSetFolder.append(ProjectFolder+"Labeled25"),DataSetName.append("Barn2")
# DataSetFolder.append(ProjectFolder+"Labeled25_10"),DataSetName.append("Barn2_10")
# DataSetFolder.append(ProjectFolder+"Labeled25_20"),DataSetName.append("Barn2_20")
# DataSetFolder.append(ProjectFolder+"Labeled25_30"),DataSetName.append("Barn2_30")
# DataSetFolder.append(ProjectFolder+"Labeled25_40"),DataSetName.append("Barn2_40")
# DataSetFolder.append(ProjectFolder+"Labeled25_50"),DataSetName.append("Barn2_50")
# DataSetFolder.append(ProjectFolder+"Labeled25_60"),DataSetName.append("Barn2_60")
# DataSetFolder.append(ProjectFolder+"Labeled25_70"),DataSetName.append("Barn2_70")
# DataSetFolder.append(ProjectFolder+"Labeled25_80"),DataSetName.append("Barn2_80")
# DataSetFolder.append(ProjectFolder+"Labeled25_90"),DataSetName.append("Barn2_90")
# DataSetFolder.append(ProjectFolder+"Labeled25_60B1"),DataSetName.append("Barn2_60B1")
# DataSetFolder.append(ProjectFolder+"Labeled25_60B2"),DataSetName.append("Barn2_60B2")
# DataSetFolder.append(ProjectFolder+"Labeled25_60B3"),DataSetName.append("Barn2_60B3")
# DataSetFolder.append(ProjectFolder+"Labeled25_60B4"),DataSetName.append("Barn2_60B4")
# DataSetFolder.append(ProjectFolder+"Labeled25_60B5"),DataSetName.append("Barn2_60B5")
# DataSetFolder.append(ProjectFolder+"Labeled25_60B6"),DataSetName.append("Barn2_60B6")
# DataSetFolder.append(ProjectFolder+"Labeled25_60B7"),DataSetName.append("Barn2_60B7")
# DataSetFolder.append(ProjectFolder+"Labeled25_60B8"),DataSetName.append("Barn2_60B8")
# DataSetFolder.append(ProjectFolder+"Labeled25_60B9"),DataSetName.append("Barn2_60B9")
# DataSetFolder.append(ProjectFolder+"Labeled25_60B10"),DataSetName.append("Barn2_60B10")
PretrainedModelName="BarnP"
FoldN=10

Vers_i=0
print("Model " + ModelType + ", pre-trained " + PretrainedModelName)

for DataSetFolder_i in range(len(DataSetFolder)):
    for WindowSize in WindowSizeList:
        ModelFileName=ModelFolder+"/"+ModelType+PretrainedModelName+"WS"+str(WindowSize)+"Fold"+str(0)+"V"+str(Vers_i)
        for FrozenLayer_i in range(FrozenLayersN):
            for Fold_i in range(FoldN+1):
                print("Training " + DataSetName[DataSetFolder_i] + ", window " + str(WindowSize) + "sec, frozen " + str(FrozenLayer_i+1) + ", fold " + str(Fold_i))
                if Fold_i==0:
                    DataFileName=DataSetFolder[DataSetFolder_i]+"/"+'FeedingBehaviour_Training'+"_WS"+str(WindowSize)+"_F"+str(Freq)+'.h5'
                else:
                    DataFileName=DataSetFolder[DataSetFolder_i]+"/"+'FeedingBehaviour_Training'+"_WS"+str(WindowSize)+"_F"+str(Freq)+"_Fold"+str(Fold_i)+'.h5'
                print(ModelFileName)
                print(DataFileName)
                f = h5py.File(DataFileName, 'r')
                ASlicedTraining = f['AccXYZ'][...]
                LabelSlicedOneHotTraining = f['LabelOneHot'][...]
                LabelSlicedTraining = f['Label'][...]
                f.close()

                TransferLearningModelFileName=ModelFileName+"_TL"+DataSetName[DataSetFolder_i]+"_L"+str(FrozenLayer_i+1)+"Fold"+str(Fold_i)+"V"+str(Vers_i)
                print(TransferLearningModelFileName)
                epochs, batch_size, verbose = 30*0+1+50*0, 32, 1

                model=tensorflow.keras.models.load_model(ModelFileName)
                for i in range(FrozenLayer_i+1):
                    model.layers[i].trainable = False
                model.compile(loss='categorical_crossentropy', optimizer='adam', metrics=["accuracy", "categorical_accuracy"])
                # print(model.summary())
                train_history = model.fit(ASlicedTraining, LabelSlicedOneHotTraining, epochs=epochs, batch_size=batch_size, verbose=verbose)
                model.save(TransferLearningModelFileName)

                print(train_history.history['loss'])
                p=train_history.history['loss']
                with open(TransferLearningModelFileName+'/train_history.csv', 'w') as f:
                    for item in p:
                        f.write("%s\n" % item)
print('\a')
```
