## Supplementary material for "CNN and transfer learning-based classification model for automated cow’s feeding behaviour recognition from accelerometer data": Code: Main_FeedingBehaviour_PrepareDatasetH5PY.py

```
# .\venv\Scripts\activate
# cd D:/CowBhave1
# python D:/CowBhave1/Main_FeedingBehaviour_PrepareDatasetH5PY.py
# https://github.com/christianversloot/machine-learning-articles/blob/main/how-to-use-h5py-and-keras-to-train-with-data-from-hdf5-files.md

import h5py
import numpy as np
from FeedingBehavior_NNlib import *
from sklearn.model_selection import KFold

ProjectFolder="D:/CowBhave1/"

DataFolder,DataSetName=[],[]
# DataFolder.append(ProjectFolder+"Labeled25"),DataSetName.append("Barn2")
DataFolder.append(ProjectFolder+"Pavlovic21/Labeled25"),DataSetName.append("BarnP")
# DataFolder.append(ProjectFolder+"Labeled25_60B1"),DataSetName.append("Barn2_60B1")
# DataFolder.append(ProjectFolder+"Labeled25_60B2"),DataSetName.append("Barn2_60B2")
# DataFolder.append(ProjectFolder+"Labeled25_60B3"),DataSetName.append("Barn2_60B3")
# DataFolder.append(ProjectFolder+"Labeled25_60B4"),DataSetName.append("Barn2_60B4")
# DataFolder.append(ProjectFolder+"Labeled25_60B5"),DataSetName.append("Barn2_60B5")
# DataFolder.append(ProjectFolder+"Labeled25_60B6"),DataSetName.append("Barn2_60B6")
# DataFolder.append(ProjectFolder+"Labeled25_60B7"),DataSetName.append("Barn2_60B7")
# DataFolder.append(ProjectFolder+"Labeled25_60B8"),DataSetName.append("Barn2_60B8")
# DataFolder.append(ProjectFolder+"Labeled25_60B9"),DataSetName.append("Barn2_60B9")
# DataFolder.append(ProjectFolder+"Labeled25_60B10"),DataSetName.append("Barn2_60B10")
# DataFolder.append(ProjectFolder+"Labeled25_5"),DataSetName.append("Barn2_5")
# DataFolder.append(ProjectFolder+"Labeled25_10"),DataSetName.append("Barn2_10")
# DataFolder.append(ProjectFolder+"Labeled25_20"),DataSetName.append("Barn2_20")
# DataFolder.append(ProjectFolder+"Labeled25_30"),DataSetName.append("Barn2_30")
# DataFolder.append(ProjectFolder+"Labeled25_40"),DataSetName.append("Barn2_40")
# DataFolder.append(ProjectFolder+"Labeled25_50"),DataSetName.append("Barn2_50")
# DataFolder.append(ProjectFolder+"Labeled25_60"),DataSetName.append("Barn2_60")
# DataFolder.append(ProjectFolder+"Labeled25_70"),DataSetName.append("Barn2_70")
# DataFolder.append(ProjectFolder+"Labeled25_80"),DataSetName.append("Barn2_80")
# DataFolder.append(ProjectFolder+"Labeled25_90"),DataSetName.append("Barn2_90")

WindowSizeList=[5,10,20,30,60,90,120,180,300] #sec
# WindowSizeList=[60] #sec
Freq=25 #Hz
FoldN=10

for DataFolder_i in range(len(DataFolder)):
    CowNoList=[]
    FileList=os.listdir(DataFolder[DataFolder_i])
    for FileName in FileList:
        if FileName.endswith(".csv") and "AccDataLabeled_Tag" in FileName:
            print(FileName)
            a=FileName.find('_Cow')
            b=FileName.find('_',a+4)
            CowNoList.append(FileName[(a+4):b])
    print(DataFolder[DataFolder_i])
    print("Cow No list")
    CowNoList=numpy.unique(CowNoList)
    print(str(len(CowNoList))+' cows: '+str(CowNoList))
    if FoldN>1:
        kf = KFold(n_splits=FoldN)
        i=1
        for train_index, test_index in kf.split(CowNoList):
            print('Fold '+str(i)+' '+str(CowNoList[train_index]))
            print(str(CowNoList[test_index]))
            i=i+1

    for WindowSize in WindowSizeList:
        WindowN=int(WindowSize*Freq)
        print("WS "+str(WindowSize))
        fTraining = h5py.File(DataFolder[DataFolder_i]+"/"+"FeedingBehaviour_Training_WS"+str(WindowSize)+"_F"+str(Freq)+'.h5', 'a')
        fValidation=h5py.File(DataFolder[DataFolder_i]+"/"+"FeedingBehaviour_Validation_WS"+str(WindowSize)+"_F"+str(Freq)+'.h5', 'a')
        i=0
        for FileName in FileList:
            if FileName.endswith(".csv") and "AccDataLabeled_Tag" in FileName:
                TagNo, CowNo, TimeStamp, Ax, Ay, Az, Label=ReadLabeledDataFiles(DataFolder[DataFolder_i],FileName,Freq)
                AxSlicedT, AySlicedT, AzSlicedT, LabelSlicedT, TagNoSlicedT, CowNoSlicedT, TimeStampSlicedT=LabeledDataSlicing(Ax, Ay, Az, Label, WindowN, 0.5, TagNo, CowNo, TimeStamp)
                AxSlicedT, AySlicedT, AzSlicedT, LabelSlicedT, CowNoSlicedT=DataBalance(AxSlicedT, AySlicedT, AzSlicedT, LabelSlicedT, CowNoSlicedT)
                AxSlicedT, AySlicedT, AzSlicedT, LabelSlicedT, CowNoSlicedT=AccRotationAugmentation(AxSlicedT, AySlicedT, AzSlicedT, LabelSlicedT, CowNoSlicedT)
                ASlicedT=numpy.dstack((AxSlicedT,AySlicedT,AzSlicedT))
                LabelSlicedT=numpy.asarray(LabelSlicedT) - 1
                LabelSlicedOneHotT=to_categorical(LabelSlicedT)
                LabelSlicedT=numpy.asarray(LabelSlicedT) + 1

                AxSlicedV, AySlicedV, AzSlicedV, LabelSlicedV, TagNoSlicedV, CowNoSlicedV, TimeStampSlicedV=LabeledDataSlicing(Ax, Ay, Az, Label, WindowN, 0.5, TagNo, CowNo, TimeStamp)
                ASlicedV=numpy.dstack((AxSlicedV,AySlicedV,AzSlicedV))
                LabelSlicedV=numpy.asarray(LabelSlicedV) - 1
                LabelSlicedOneHotV=to_categorical(LabelSlicedV)
                LabelSlicedV=numpy.asarray(LabelSlicedV) + 1

                if i==0:
                    fTraining.create_dataset("AccXYZ", data=ASlicedT, chunks=True, maxshape=(None,ASlicedT.shape[1],ASlicedT.shape[2]))
                    fTraining.create_dataset("LabelOneHot", data=LabelSlicedOneHotT, chunks=True, maxshape=(None,LabelSlicedOneHotT.shape[1]))
                    fTraining.create_dataset("Label", data=LabelSlicedT, chunks=True, maxshape=(None,))

                    fValidation.create_dataset("AccXYZ", data=ASlicedV, chunks=True, maxshape=(None,ASlicedV.shape[1],ASlicedV.shape[2]))
                    fValidation.create_dataset("LabelOneHot", data=LabelSlicedOneHotV, chunks=True, maxshape=(None,LabelSlicedOneHotV.shape[1]))
                    fValidation.create_dataset("Label", data=LabelSlicedV, chunks=True, maxshape=(None,))
                    i=1
                else:
                    fTraining["AccXYZ"].resize((fTraining["AccXYZ"].shape[0] + ASlicedT.shape[0]), axis = 0)
                    fTraining["AccXYZ"][-ASlicedT.shape[0]:] = ASlicedT
                    fTraining["LabelOneHot"].resize((fTraining["LabelOneHot"].shape[0] + LabelSlicedOneHotT.shape[0]), axis = 0)
                    fTraining["LabelOneHot"][-LabelSlicedOneHotT.shape[0]:] = LabelSlicedOneHotT
                    fTraining["Label"].resize((fTraining["Label"].shape[0] + LabelSlicedT.shape[0]), axis = 0)
                    fTraining["Label"][-LabelSlicedT.shape[0]:] = LabelSlicedT

                    fValidation["AccXYZ"].resize((fValidation["AccXYZ"].shape[0] + ASlicedV.shape[0]), axis = 0)
                    fValidation["AccXYZ"][-ASlicedV.shape[0]:] = ASlicedV
                    fValidation["LabelOneHot"].resize((fValidation["LabelOneHot"].shape[0] + LabelSlicedOneHotV.shape[0]), axis = 0)
                    fValidation["LabelOneHot"][-LabelSlicedOneHotV.shape[0]:] = LabelSlicedOneHotV
                    fValidation["Label"].resize((fValidation["Label"].shape[0] + LabelSlicedV.shape[0]), axis = 0)
                    fValidation["Label"][-LabelSlicedV.shape[0]:] = LabelSlicedV
        fTraining.close()
        fValidation.close()

        if FoldN<=1:
            continue

        kf = KFold(n_splits=FoldN)
        #Training
        print("Training folds")
        Fold_i=0
        for train_index, test_index in kf.split(CowNoList):
            Fold_i=Fold_i+1
            jt,jv=0,0
            DataFileName=DataFolder[DataFolder_i]+"/"+"FeedingBehaviour_Training_WS"+str(WindowSize)+"_F"+str(Freq)+"_Fold"+str(Fold_i)+'.h5'
            fTrainingFold = h5py.File(DataFileName, 'a')
            DataFileName=DataFolder[DataFolder_i]+"/"+"FeedingBehaviour_Validation_WS"+str(WindowSize)+"_F"+str(Freq)+"_Fold"+str(Fold_i)+'.h5'
            fValidationFold=h5py.File(DataFileName, 'a')
            print("Interval length " + str(WindowSize) + "sec, " + "Fold "+str(Fold_i)+", training="+str(CowNoList[train_index])+", validation="+str(CowNoList[test_index]))
            
            for FileName in FileList:
                if FileName.endswith(".csv") and "AccDataLabeled_Tag" in FileName:
                    TagNo, CowNo, TimeStamp, Ax, Ay, Az, Label=ReadLabeledDataFiles(DataFolder[DataFolder_i],FileName,Freq)
                    if CowNo[0] in CowNoList[train_index]:
                        AxSlicedT, AySlicedT, AzSlicedT, LabelSlicedT, TagNoSlicedT, CowNoSlicedT, TimeStampSlicedT=LabeledDataSlicing(Ax, Ay, Az, Label, WindowN, 0.5, TagNo, CowNo, TimeStamp)
                        AxSlicedT, AySlicedT, AzSlicedT, LabelSlicedT, CowNoSlicedT=DataBalance(AxSlicedT, AySlicedT, AzSlicedT, LabelSlicedT, CowNoSlicedT)
                        AxSlicedT, AySlicedT, AzSlicedT, LabelSlicedT, CowNoSlicedT=AccRotationAugmentation(AxSlicedT, AySlicedT, AzSlicedT, LabelSlicedT, CowNoSlicedT)
                        ASlicedT=numpy.dstack((AxSlicedT,AySlicedT,AzSlicedT))
                        LabelSlicedT=numpy.asarray(LabelSlicedT) - 1
                        LabelSlicedOneHotT=to_categorical(LabelSlicedT)
                        LabelSlicedT=numpy.asarray(LabelSlicedT) + 1
                        if jt==0:
                            fTrainingFold.create_dataset("AccXYZ", data=ASlicedT, chunks=True, maxshape=(None,ASlicedT.shape[1],ASlicedT.shape[2]))
                            fTrainingFold.create_dataset("LabelOneHot", data=LabelSlicedOneHotT, chunks=True, maxshape=(None,LabelSlicedOneHotT.shape[1]))
                            fTrainingFold.create_dataset("Label", data=LabelSlicedT, chunks=True, maxshape=(None,))
                            jt=1
                        else:
                            fTrainingFold["AccXYZ"].resize((fTrainingFold["AccXYZ"].shape[0] + ASlicedT.shape[0]), axis = 0)
                            fTrainingFold["AccXYZ"][-ASlicedT.shape[0]:] = ASlicedT
                            fTrainingFold["LabelOneHot"].resize((fTrainingFold["LabelOneHot"].shape[0] + LabelSlicedOneHotT.shape[0]), axis = 0)
                            fTrainingFold["LabelOneHot"][-LabelSlicedOneHotT.shape[0]:] = LabelSlicedOneHotT
                            fTrainingFold["Label"].resize((fTrainingFold["Label"].shape[0] + LabelSlicedT.shape[0]), axis = 0)
                            fTrainingFold["Label"][-LabelSlicedT.shape[0]:] = LabelSlicedT
                    
                    elif CowNo[0] in CowNoList[test_index]:
                        AxSlicedV, AySlicedV, AzSlicedV, LabelSlicedV, TagNoSlicedV, CowNoSlicedV, TimeStampSlicedV=LabeledDataSlicing(Ax, Ay, Az, Label, WindowN, 0.5, TagNo, CowNo, TimeStamp)
                        ASlicedV=numpy.dstack((AxSlicedV,AySlicedV,AzSlicedV))
                        LabelSlicedV=numpy.asarray(LabelSlicedV) - 1
                        LabelSlicedOneHotV=to_categorical(LabelSlicedV)
                        LabelSlicedV=numpy.asarray(LabelSlicedV) + 1
                        if jv==0:
                            fValidationFold.create_dataset("AccXYZ", data=ASlicedV, chunks=True, maxshape=(None,ASlicedV.shape[1],ASlicedV.shape[2]))
                            fValidationFold.create_dataset("LabelOneHot", data=LabelSlicedOneHotV, chunks=True, maxshape=(None,LabelSlicedOneHotV.shape[1]))
                            fValidationFold.create_dataset("Label", data=LabelSlicedV, chunks=True, maxshape=(None,))
                            jv=1
                        else:
                            fValidationFold["AccXYZ"].resize((fValidationFold["AccXYZ"].shape[0] + ASlicedV.shape[0]), axis = 0)
                            fValidationFold["AccXYZ"][-ASlicedV.shape[0]:] = ASlicedV
                            fValidationFold["LabelOneHot"].resize((fValidationFold["LabelOneHot"].shape[0] + LabelSlicedOneHotV.shape[0]), axis = 0)
                            fValidationFold["LabelOneHot"][-LabelSlicedOneHotV.shape[0]:] = LabelSlicedOneHotV
                            fValidationFold["Label"].resize((fValidationFold["Label"].shape[0] + LabelSlicedV.shape[0]), axis = 0)
                            fValidationFold["Label"][-LabelSlicedV.shape[0]:] = LabelSlicedV

            fTrainingFold.close()
            fValidationFold.close()
        
print('\a')
```
